# The splicing factor U2AF1 contributes to cancer progression through a non-canonical role in translation regulation

**DOI:** 10.1101/371492

**Authors:** Murali Palangat, Dimitrios Anastasakis, Fei Liang, Katherine E. Lindblad, Robert Bradley, Christopher S. Hourigan, Markus Hafner, Daniel R. Larson

## Abstract

Somatic mutations in the genes encoding components of the spliceosome occur frequently in human neoplasms, including myeloid dysplasias and leukemias and less often in solid tumors. One of the affected factors, U2AF1, is involved in splice site selection, and the most common change, S34F, alters a conserved nucleic acid binding domain, recognition of the 3’-splice site, and alternative splicing of many mRNAs. However, the role that this mutation plays in oncogenesis is still unknown. Here, we have uncovered a non-canonical function of U2AF1, showing that it binds mature mRNA in the cytoplasm and negatively regulates mRNA translation. This splicing-independent role of U2AF1 is altered by the S34F mutation, and polysomal profiling indicates the mutation affects translation of hundreds of mRNA. One functional consequence is increased synthesis of the secreted chemokine interleukin 8 which contributes to metastasis, inflammation, and cancer progression in mice and humans.

---

Somatic mutations affecting the genes that encode core components of the splicing machinery, first discovered in blood malignancies^1^, have now been observed in nearly every cancer type^2–4^. These somatic mutations are largely present in factors including SF3B1, SRSF2, ZRSR2 and U2AF1, that are involved in selection of branch-point, exonic sequences, and 3’-splice sites. Spliceosome mutations result in hundreds of changes in alternative splicing, which vary among tissues and patients^5^. The spectrum of mutations, the stoichiometry in tumors, and the clinical outcomes point toward splicing factor mutations as oncogenic events^4^. However, there is little consensus on the molecular mechanism of action in human carcinogenesis.

The splicing factor U2AF, a heterodimer of U2AF1 and U2AF2, is responsible for the recognition and binding to the 3’splice site (U2AF1 to AG nucleotides and U2AF2 to the polypyrimidine tract), a critical initial step in the assembly of the spliceosome^6–8^. A recurrent hotspot mutation in U2AF1 (S34F) in its first Zn-finger domain^1,9,10^ critical for RNA binding activity^11^, alters RNA binding specificity^12–15^ and splicing kinetics^16^, resulting in a wide variety of splicing outcomes, most frequently cassette exon inclusion^1,3,10,15,17–19^. In all cases, the tumor cells retain a wild type allele of U2AF1^3^. Most of the splicing changes observed in human tissue can be recapitulated in a <u>H</u>uman <u>B</u>ronchial <u>E</u>pithelial <u>C</u>ell (HBEC) line carrying the heterozygous U2AF1-S34F mutation^12^. However, mutant HBECs do not exhibit any visible morphological or altered growth phenotypes^12^, indicating the mutation is not a strong driver mutation. Although the S34F mutation mediates subtle splicing changes, these isoform shifts have not been definitively linked to a phenotype, leading some investigators to propose nuclear RNA processing defects such as alterations in 3’ UTRs^20^ or an increase in R-loops^21^.

Here, we show that U2AF1 functions as a translational repressor in the cytoplasm. U2AF1 interacts with hundreds of spliced, poly-adenylated mRNA in the cytoplasm, many of which are mTOR-regulated and code for proteins involved in translation initiation and growth. Through RIP-seq and polysome profiling, we show that the cancer-associated S34F mutation results in loss of binding and translational de-repression. We further provide evidence in support of a functional role for this non-canonical pathway through the translational regulation of the chemokine IL8. IL8 is translationally up-regulated in the mutant background, increases epithelial to mesenchymal transition in culture, enhances the inflammatory response in mice, and blocking IL8 with neutralizing antibody reduces tumor burden. Moreover, elevated IL8 levels in human bone marrow correlate with acute myeloid leukemia. Thus, U2AF1 S34F has the potential to play both cell autonomous and non-autonomous roles in cancer progression, and the mechanism described here suggests a possible therapeutic intervention.

## Results

### U2AF1-S34F mutation confers altered viability and sustained inflammatory cytokine secretion after DNA damage

To investigate the mechanism of the S34F mutation in oncogenesis, we used four isogenic HBEC cell lines created with gene-editing^12^: 1) wt/wt, the parent cell line; 2) wt/S34F, the heterozygous condition similar to that observed in patients; 3) wt/-, a line with one allele carrying a frame-shift mutation; and 4) wt/S34F-, a line derived from the wt/S34F line where the mutant allele is frameshifted, resulting in removal of the mutant U2AF1-S34F protein. Using RNA-sequencing we observed modest changes (1.5- to 2.0-fold increase or decrease) in gene expression in the wt/S34F mutant cells and altered splicing outcomes (Extended Data Fig. 1a-c). Genes displaying changes in RNA levels were enriched for those associated with DNA metabolism, (Extended Data Table 1), suggesting either changes in cell cycle progression or DNA damage.

Although we did not observe change in growth rates^12^, we did find that the S34F mutation conferred altered outcomes upon DNA damage. When irradiated with a high dose of X-rays (20 Gy), the S34F mutation conferred substantial growth advantage over a 6-day period in a cell proliferation assay (Fig. 1a), and this phenotype could be successfully rescued with the frame-shift mutation, indicating the S34F mutation resulted in gain of function. A fraction of wt/S34F cells survived irradiation and stained positive for the senescence marker β-galactosidase and remained in a senescent state in culture for > 30 days, whereas the wt/wt and wt/− mutant cells died 10-12 days post irradiation (Extended Data Fig. 1d). Thus, U2AF1 S34F provides resistance to clinically relevant doses of X-rays.

**Figure 1.**
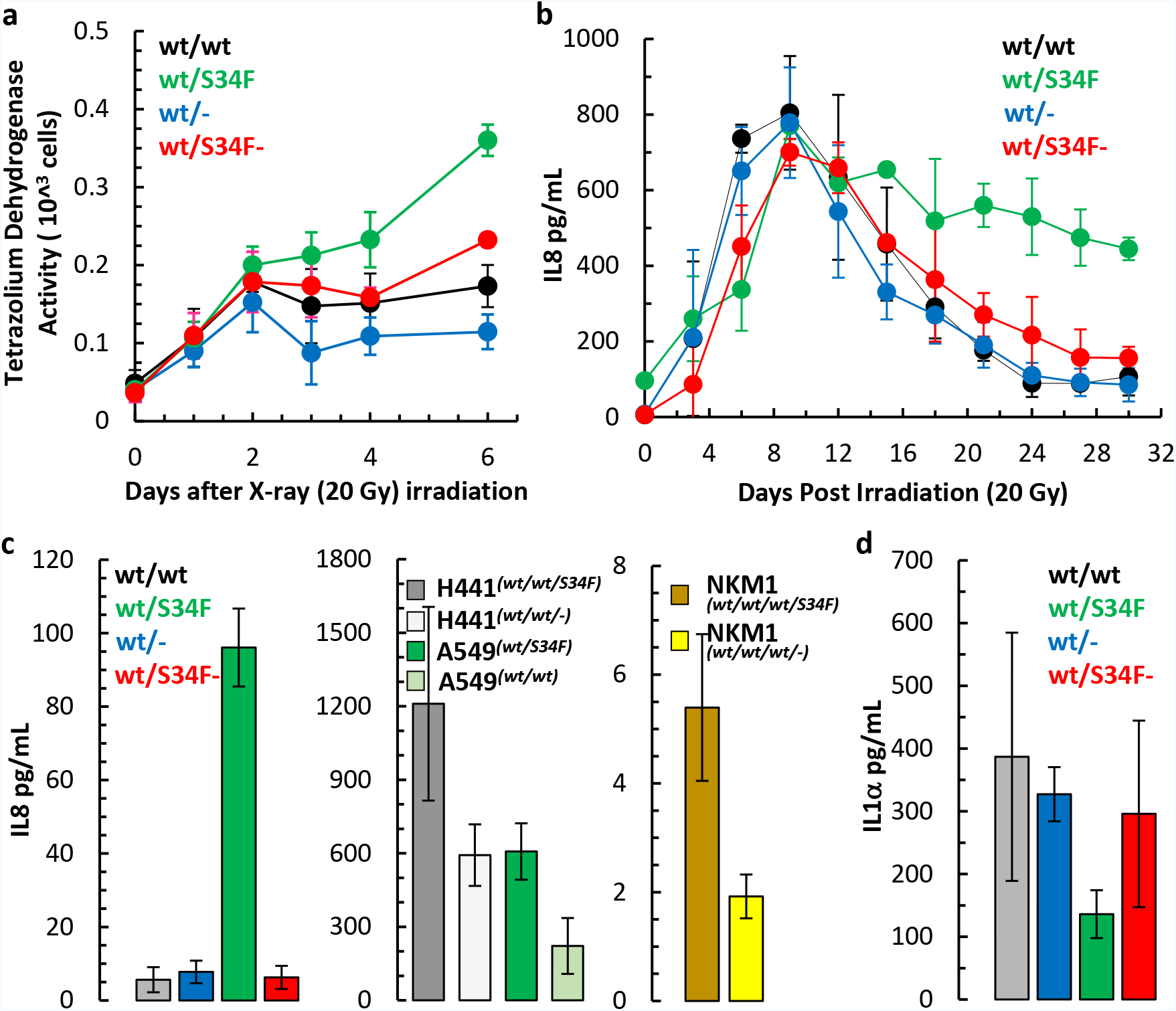
U2AF1 S34F mutant cells show altered DNA damage response and inflammatory cytokine secretion. a. Altered growth in wt/wt, wt/S34F, wt/− and wt/S34F-mutant cells after irradiation with x-rays (20 Gy) measured using a cell proliferation assay at indicated times (average of 3 independent experiments +/− s.d.). b. Cytokine levels measured by ELISA in media collected at indicated times from wt/wt, wt/S34F, wt/− and wt/S34F-mutant cells after irradiation with x-rays (20 Gy). IL8 secreted into media is presented as pg/mL (average of 3 independent experiments +/− s.d.). c. ELISA of steady state secretion of IL8 into media collected after 6 days in culture of HBEC (wt/wt, wt/S34F, wt/− and wt/S34F-, (wt/wt *vs* wt/S34F p = 0.0003; wt/wt *vs* wt/S34F− p = 0.8533), human lung tumor cells (H441 and A549, H441^S34F^ *vs* H441^S34F-^ p = 0.0614; A549^S34F^ *vs* A549^S34F-^ p = 0.0281) and human myeloid leukemia cell line (NKM1, NKM1^S34F^ *vs* NKM1^S34F-^ p = 0.0251) and with their respective S34F mutant allele knockout cell lines (average of 3 biological replicates +/− s.d.). d. ELISA of steady state secretion of IL1-α into media collected after 6 days in culture of wt/wt, wt/S34F, wt/− and wt/S34F-mutant cells (average of 3 biological replicates +/− s.d. wt/wt *vs* wt/S34F p = 0.3446; wt/wt *vs* wt/S34F− p = 0.6601). P values determined by two sided T-Test.

These results bear resemblance to the phenomenon of oncogene-induced senescence, where non-proliferating cells contribute to oncogenesis through secretion of factors into the surrounding tissue^22–25^. We investigated if these cells secreted analytes characteristic of senescent cells, and tested by ELISA for a panel of 10 cytokines or chemokines (IL1α, IL1β, IL2, IL4, IL6, IL8, IL10, IFNγ & TNfα in media collected post irradiation over a 30-day period. Two cytokines – IL8 and IL1α – showed altered secretion patterns. Strikingly, the senescent wt/S34F mutant cells exhibited sustained secretion of IL8 for over 30 days, and this phenotype could be rescued by frame-shifting the mutant allele (Fig. 1b). Even at steady state (no irradiation), the wt/S34F mutant cells secreted elevated levels of IL8 compared to wt/wt cells (17-fold increase), and this phenomenon was rescued in the wt/S34F-line (Fig. 1c). We confirmed this observation in human lung cancer cell lines A549 and H441 and the myeloid leukemia cell line NKM1 which were either gene-edited to create the S34F mutation or correct a naturally-occurring U2AF1-S34F mutation: each cell line secreted elevated levels of IL8 in a mutant-dependent manner (Fig. 1c). In contrast, the wt/S34F HBEC cells showed decreased secretion of IL1α relative to wt/wt cells before and after irradiation, an effect which was also rescued in wt/S34F-cells (Fig. 1d; Extended Data Figure 1e). In summary, U2AF1-S34F cells show radiation resistance and altered secretion of inflammatory cytokines even before treatment with X-rays. Because of the role of inflammatory cytokines in cancer^26,27^, we chose to study this phenotype further.

### U2AF1 is present in the cytoplasm and binds mature mRNA

Surprisingly, we observed no changes in IL8 splicing by RNA-seq^12^. Moreover, we did not observe any significant change in IL8 expression between wt/wt and wt/S34F mutant cells as measured by single-molecule RNA-FISH and gene expression (RT-qPCR) analyses (Fig. 2a & Extended Data Fig 2a). Previous studies have shown IL8 expression is regulated both at the level of mRNA stability and translational control^28–30^. Since there was no change in IL8 mRNA levels, the elevated secretion of IL8 in wt/S34F cells is controlled either at the level of translation or trafficking/secretion. We therefore sought to determine if U2AF1 could be playing a role in regulation of this message independent of splicing.

**Figure 2.**
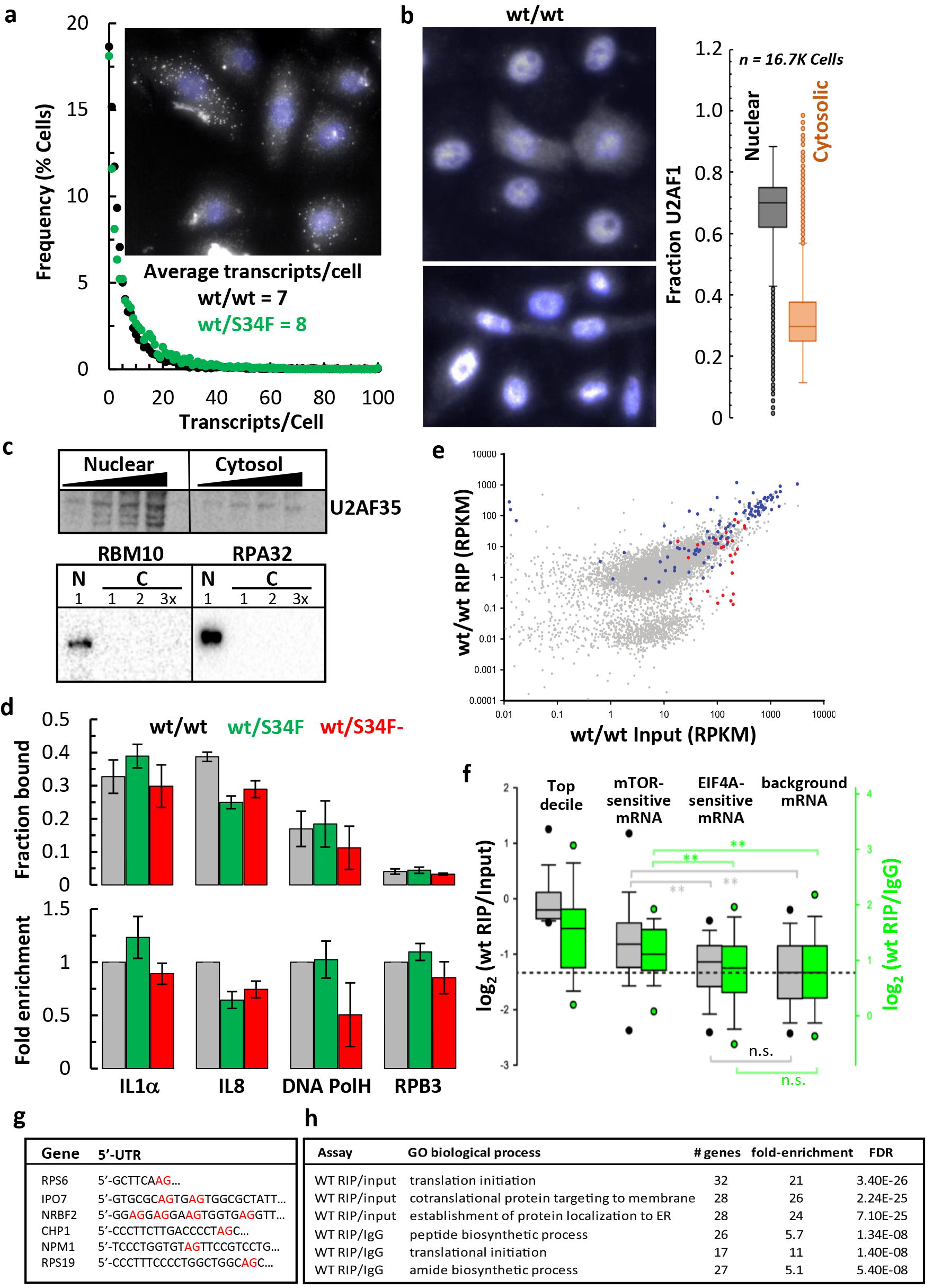
U2AF1 binds RNA in the cytoplasm. a. Single molecule FISH analysis of IL8 mRNA expression in wt/wt, wt/S34F mutant cells plotted as frequency histogram of IL8 mRNA per cell. Inset: single molecule RNAFISH image (Gray) showing heterogeneity in IL8 RNA expression and nuclei counter stained with DAPI (Blue) in wt/wt cells. b. Quantitative immunofluorescence detection of cytoplasmic U2AF1 (Gray) in PFA fixed cells showing heterogeneity in U2AF1 distribution in the cytoplasm and nuclei counter stained with DAPI (Blue). Box plot (min, max, 1^st^ & 3^rd^ quartiles and median) showing the fraction of U2AF1 in the nucleus and cytoplasm of individual wt/wt cells. c. Western blot analysis showing presence of U2AF1 in the cytosolic fraction. RBM10 and RPA32 (increasing amounts) served as controls for any nuclear protein contamination in the cytoplasmic fraction. d. U2AF1 inteacts with mature IL8 mRNA in the cytoplasm. RNA isolated from immunoprecipitates of cytoplasmic U2AF1 (b) was analyzed by RT-qPCR for association of IL8, IL1α, DNA polH and RPB3 mRNA. Fraction of IL1α, IL8, DNA polH and RPB3 mRNA bound to U2AF1 (top) and their relative enrichment to U2AF1 in mutant cells (bottom). e. Transcriptome-wide analysis of WT U2AF1 bound cytoplasmic mature polyadenylated mRNA by RIP-seq. Scatter plot of enrichment of U2AF1 bound RNA (gray), intron-less Histone RNA (red) and mTOR regulated RNA (blue). f. Box plot analysis of mRNA enriched in RIP over input or control IgG in the top decile of RIP, mTOR sensitive mRNA, EIF4A sensitive mRNA or background mRNA. g. 3’-Splice site like “AG” sequence and polypyrimidine rich TOP motifs in 5’-UTRs of mTOR regulated mRNAs bound by cytoplasmic U2AF1. h. Gene Ontology terms of wild type U2AF1 enriched cytoplasmic mRNA.

We first asked whether U2AF1 interacts with IL8 mRNA. Fluorescently labeled U2AF1 has been shown to shuttle between the nucleus and cytoplasm in plant cells^31^ and using a heterokaryon assay in human cells^32^. Also, splicing factors have been implicated in translational control^33–35^, leading us to ask whether there exists a cytoplasmic population of U2AF1 in human cells. We observed localisation of U2AF1 in the cytoplasm of wt/wt cells by immunostaining with ~30% localised in the cytoplasm (Fig. 2b), and we could immunoprecipitate U2AF1 from the cytoplasmic fraction with minimal nuclear contamination (Fig. 2c, Extended Data Fig. 2b & c). We next tested for the presence of mature spliced, poly-adenylated mRNA in the cytoplasmic U2AF1-immunoprecipitates and detected the presence of mRNA for IL1α, IL8 and DNA pol H in substantial quantities, but very little of RPB3 mRNA (Fig. 2d, Extended Data Fig 2d), suggesting preferential binding of U2AF1 to certain mRNAs and not others. In the wt/S34F mutant cells, binding of U2AF1 to IL8 mRNA was significantly lower (Fig. 2d; Extended Data Fig 2d; note: wt/S34F cells are heterozygous and the antibody recognizes both wt and S34F forms of U2AF1).

We next sought to generate a comprehensive transcriptome-wide view of U2AF1 interactions with RNA in the cytoplasm. RNA-IP under native conditions was performed on cytosolic extracts with anti-U2AF1 followed by sequencing (RIP-seq) of associated RNAs. Strikingly, we observed that mRNAs whose translation is regulated by mTOR are highly enriched in the U2AF1 RIP in wt/wt cells (Fig. 2e, blue circles, Extended Data Table 2). In fact, in a rank ordered-list of RIP/input for all messages exceeding a threshold for expression (1,117 transcripts, RPKM > 1 in all fractions and samples; Extended Data Table 3) some of the top hits are mTOR-regulated mRNA (RPS6, IPO7, NPM1, RPS19). Moreover, mTOR-regulated mRNAs as a class^36^ show greater enrichment in the U2AF1 IP than either the total pool of transcripts or another class of translationally-regulated transcripts (EIF4A-senstive mRNA^37,38^) (Fig. 2f). Since mTOR-regulation is conferred by the presence of a terminal oligo-pyrimidine (TOP) motif in the 5’-UTR^36,39^ we visually inspected the 5’ UTRs of some of these messages determined experimentally through previous work^40^. We observed a sequence resembling the TOP motif often located adjacent to an AG dinucleotide recognized by U2AF1, resulting in a sequence which is similar to a canonical 3’ ss^41^ (Fig. 2g). IL8 did not meet our RIP-seq threshold criteria due to low expression levels but is reported to be sensitive to mTOR inhibitors^28,29^. Finally, we took the top quintile of hits from lists ranked either by RIP/input or RIP/IgG and analyzed the GO terms. In both rankings the enriched transcripts are those involved in translational initiation and peptide biosynthesis (Fig. 2h). Thus, wt U2AF1 specifically binds mRNA in the cytosol, and many of these messages are mTOR regulated and/or code for proteins involved in translation.

### A non-canonical function for U2AF1 as a translational repressor

The decreased binding of U2AF1 to IL8 mRNA in wt/S34F cells, coupled with increased levels of IL8 in the media, led us to consider that U2AF1 might function as a translational repressor. To test this hypothesis we performed polysome profiling. We isolated polysomes from wt/wt, wt/S34F and wt/S34F-mutant cells (Fig. 3a, b; Extended Data Fig 3a) and determined fractional amount of mRNA relative to monosomes. mRNAs of sufficient length with high initiation rates contain multiple ribosomes (polysomes) and appear in the heavy fractions of a sucrose gradient, while mRNAs with low initiation rates appear in fractions containing small polysomes or 80s monosomes^42^. In addition to IL8 and IL1α, we tested the mRNAs of NPM1 and PABPC1, two mTOR-regulated RNAs which were either enriched or not enriched as measured by U2AF1 RIP-seq (Extended data Table 3). NPM1 and IL8 show loss of binding in the wt/S34F cells and were enriched in the heavy polysome fractions (Fig. 3b). PABPC1 mRNA shows no change (Fig 3b). The enhanced translational efficiency observed for NPM1 and IL8 mRNA by polysome analysis in mutant cells was also reflected in the steady state protein levels (Fig. 3c). We also tested steady state protein levels for several other genes: IPO7 (reduced binding in wt/S34F) and TSC2 (mTOR-sensitive but not detected above threshold in RIP) showed higher levels of protein; RUNX1 and TRIM28 (no change or small gain in binding) showed no change or decreased translation respectively (Fig. 3c).

**Figure 3.**
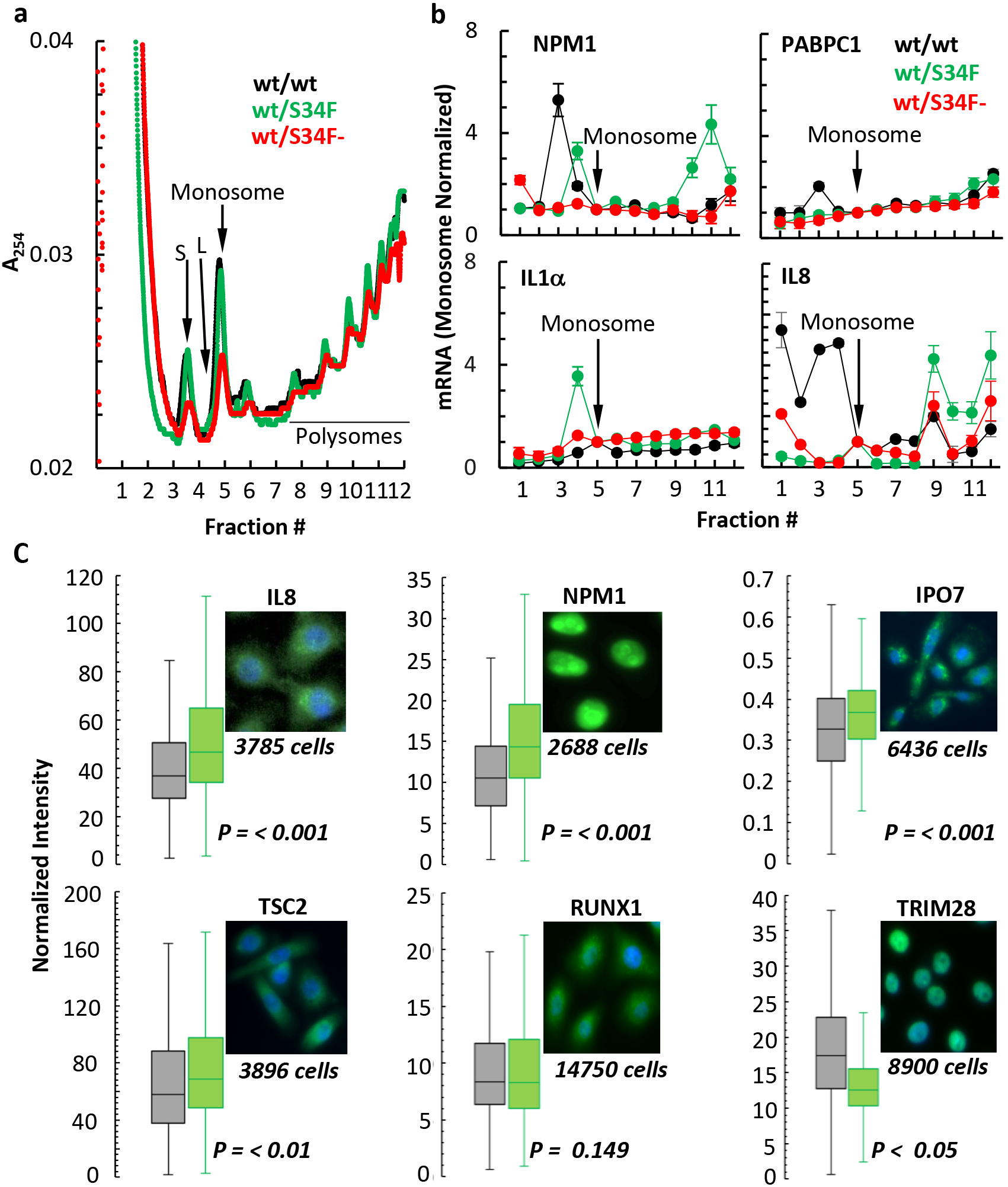
U2AF1 functions as a translational repressor in the cytoplasm. a. 10-50% linear sucrose gradient profile of polysomes isolated from wt/wt, wt/S34F and wt/S34F-mutant cells. The positions of the small (S) and large (L) ribosomal subunits, monosomes and polysomes are indicated. b. Monosome and polysome fractions (3 biological replicates) were subjected to semi-quantitative RT-PCR for indicated genes, products fractionated on a 1.5% agarose gel and quantified. The mRNA in individual fractions normalized to the total in all fractions and the ratio of mRNA to monosome in the individual fractions plotted (average of three independent experiments +/− SE). c. Box plot analysis of normalized integrated fluorescence intensity (min, max, 1^st^ & 3^rd^ quartiles and median) after immunostaining PFA fixed wt/wt (black) and wt/S34F (green) cells for IL8, NPM1, IPO7, TSC2, RUNX1 and TRIM28. P values determined by two sided T-Test.

We next sought to comprehensively test for a direct repressive role of U2AF1 in translation which might be altered by the S34F mutation by sequencing the mRNA isolated from polysome fractions (Extended Data Table 4). Individual polysome profiles are shown in Fig. 4a. NPM1, IL8, and IPO7 translational changes in wt/S34F cells were re-capitulated in the genome-wide assay, while PABPC1 showed no change in translational efficiency with mutation status. The cytoplasmic binding targets of U2AF1 from RIP-seq were then rank-ordered by loss of binding which occurred in wt/S34F cells and divided into deciles (n^~^ 112 transcripts/decile, Fig. 4b). The average polysome profiles from the top and bottom deciles demonstrate that the transcripts which showed the greatest change in binding (decile 1) also showed a polysome profile which is characteristic of more efficient translation (Fig. 4c). In fact, when looking at a single ratio of polysome to monosomes over all deciles, we observe a systematic trend where loss of binding as measured by RIP results in an increase in polysome/monosme ratio, with the top two deciles showing the most pronounced effect (Fig. 4d, green). This effect is not observed in wt/wt cells (gray) and is rescued in wt/S34F-cells (red) (Fig. 4d). Supporting this result, when transcripts are rank ordered by enrichment in the U2AF1 RIP in wt/wt cells, there is a systematic trend toward less binding resulting in greater translation (Fig. 4e, f). Thus, in independent measurements of RIP-seq and POLY-seq, U2AF1-bound transcripts show lower translation.

**Figure 4.**
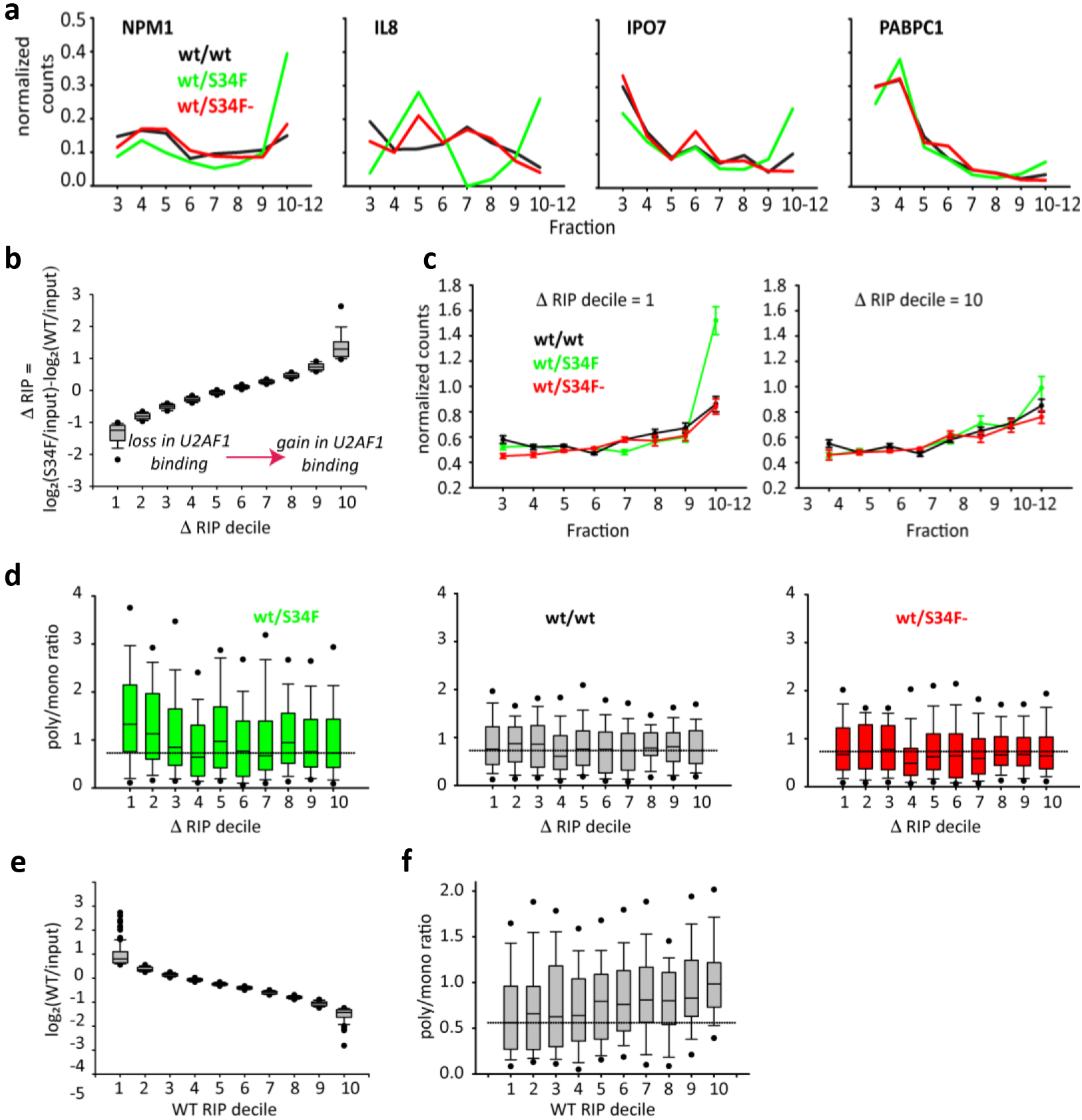
U2AF1 directly represses translation of hundreds of genes. a. Polysome profiles of indicated genes showing their enhanced distribution in heavy polysomes from wt/S34F cells. Normalization to unit area for each sample. b. Rank order of S34F mutant associated change in U2AF1 binding in RIP samples and binned into deciles (^~^112 transcripts/decile). Bootstrap error bars. c. Altered translation (sequence counts) of all mRNAs in the top (left plot) and bottom (right plot) deciles in panel b. d. Change in translation (polysomes to monosome ratio) of all mRNAs in each decile (panel b) from wt/S34F (green), wt/wt (gray) and wt.S34F- (red) cells. e. Rank order of U2AF1 associated mRNA from wt/wt cells relative to input and binned into deciles as in panel b. f. Polysome to monosome ratio of mRNA in all deciles from panel e showing reduced binding by U2AF1 to mRNA in wt/wt cells does not correlate with enhanced translation.

Loss of U2AF1 binding in wt/S34F cells is predictive of gain in translation efficiency on hundreds of messages, suggesting a functional role for U2AF1 in translational repression which is abrogated by the S34F mutation. The targets of this translational repression pathway in wt/wt cells are enriched in messages which themselves code for translation machinery (Fig. 2h). However, the most substantial *changes* in binding and translation with the S34F mutation occur for a subset of the translational machinery, in the GO category of ‘intracellular protein transport’ (Extended Data Fig. 4).

### U2AF1 regulates translation of IL8 via its 5’-UTR

Because we identified a role for U2AF1 in translation repression, we analyzed 5’ UTRs for the enrichment of the U2AF1 motif and found it to be pervasive, with 4608 out of 6441 experimentally determined 5’ UTRs containing the motif (FIMO, p<0.01). We also identified a number of 3’-ss like sequences in the IL8 5’ UTR. To test whether these motifs could be functioning as *cis*-acting U2AF1 binding sites, we made a series of IL8-T2A-GFP reporters (Fig. 5a) either with the wildtype 5’-UTR of IL8 or single point mutant in 3’-ss like sequences. This reporter contains no introns, and the GFP is not secreted, thereby testing the translational control mechanism in a splicing- and secretion-independent manner. We found that a point mutant in the canonical 3’-ss nearest the 5’ end (changing the trinucleotide sequence from “CAG” to “CAT”) showed translation de-repression. Wt/wt and wt/S34F mutant cells carrying either the wild type or mutant IL8 reporter had similar levels of RNA expression (Fig 5b), and FACS analysis revealed that the wt/S34F mutant cells showed an increase (^~^1.4 fold) in the fraction of the cells expressing GFP (Fig. 5c-e, Extended Data Fig. 5), consistent with our earlier findings (Fig. 3c; Extended Data Fig. 3c). Strikingly, we observed a 5- or 10-fold increase for the 5’ UTR mutant IL8 reporter in cells that were either wild type or S34F mutant for U2AF1 respectively (Fig. 5c-e), indicating that a point mutation in the U2AF1 motif in the IL8 5’-UTR was sufficient to overcome translational repression. Similarly, the mutated 5’-UTR significantly compromised U2AF1 binding in both wt/wt and wt/S34F cells (Fig. 5f).

**Figure 5.**
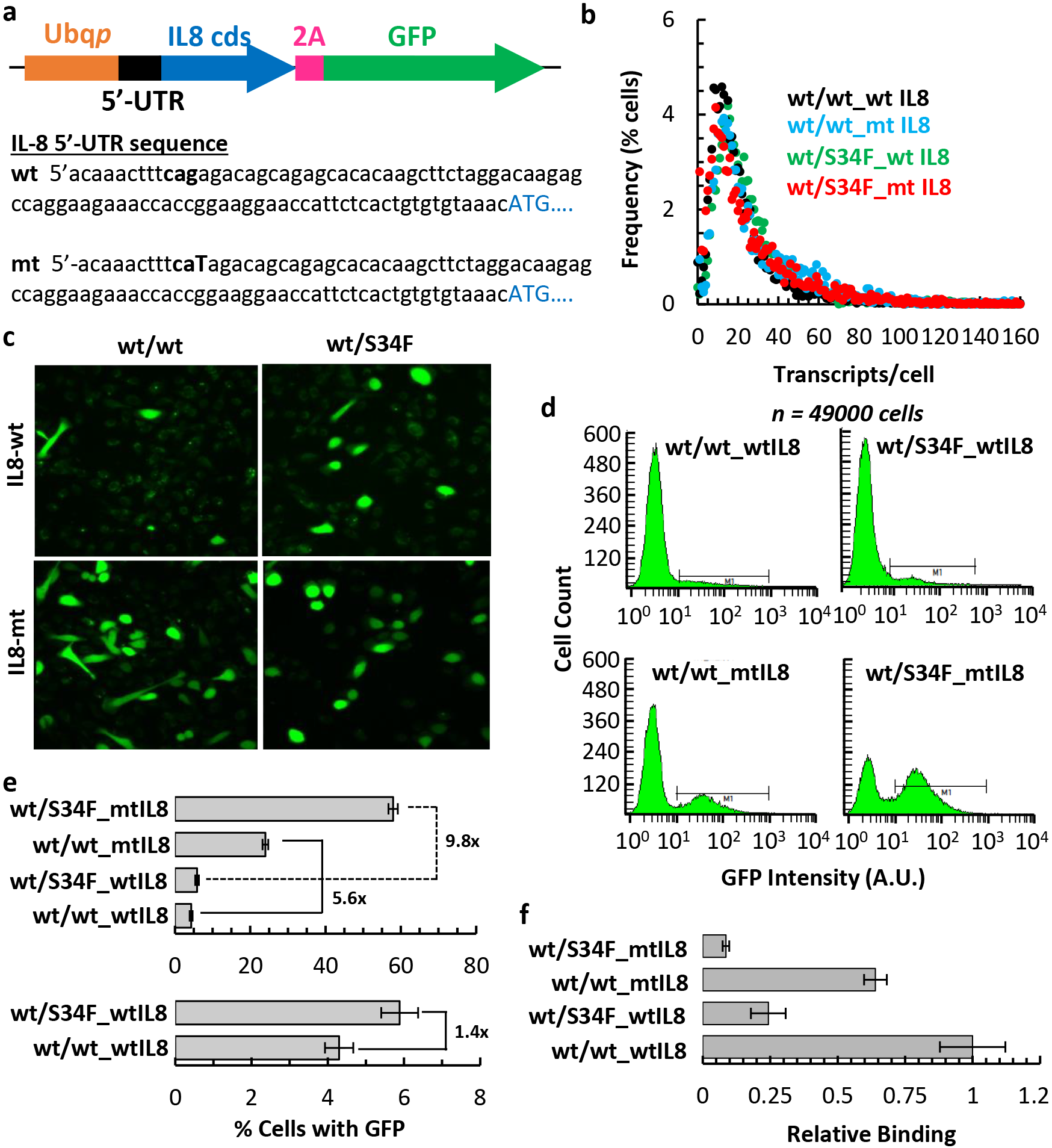
U2AF1 controls translation of IL8 via its 5’-UTR. a. Schematic representation of the IL8-2A-GFP reporter construct. The wild type IL8-5’-UTR sequence with the putative 3’-splice site like sequence depicted in bold. The mutated base within the 3’-splice site in the mutant sequence is capitalized. b. Quantitative smFISh analysis of reporter gene expression using probes specific to GFP and represented as a frequency histogram of the number of transcripts per cell. c. Confocal live cell image of GFP expression from the wt- or mt-IL8-2A-GFP reporter integrated into wt/wt or wt/S34F mutant cells. d. FACS of wt/wt or wt/S34F cells expressing wt- or mt-5’-UTR IL8-2A-GFP reporter gated above auto fluorescence from wt/wt or wt/S34F mutant control cells respectively. e. Percent cells expressing GFP from the wt- or mt-5’UTR IL8-2A-GFP reporter as a function of wild type or S34F U2AF1 status (mean of 3 biological replicates +/- SD). f. Association of wt- or mt-IL8-2A-GFP mRNA to U2AF1 in vivo as a function of wild type or S34F U2AF1 status (mean of 3 biological replicates +/- SD).

Overall, our data indicate that U2AF1 represses translation in the cytosol, that this repression is relieved in the presence of the S34F mutation on hundreds of messages, and that repression is mediated for IL8 by a *cis*-acting sequence in the 5’ UTR.

### Elevated IL8 is associated with multiple measures of cancer progression

We next asked whether this change in IL8 translation was sufficient to mediate phenotypic consequences. We carried out experiments on epithelial to mesenchymal transition (EMT), inflammatory response, and tumor burden during cancer progression. EMT is a critical step in metastasis^43–47^, known to be induced by IL8^43–45^, and is characterized by the induction of fibronectin and repression of E-Cadherin expression in epithelial cells^46,47^. Purified recombinant IL8 by itself elicited EMT in MCF-7 cells (Fig. 6a, b), as determined from the emergence of a fibronectin-high and E-cadherin-low population in tissue culture, as observed previously. Conditioned media from the wt/S34F mutant cells also induced the expression of fibronectin and repression of E-cadherin, but the conditioned media from wt/wt cells did not (Fig. 6a, b). These data indicate that the media from wt/S34F cells was biologically active and was sufficient to induce EMT-related gene expression changes.

**Figure 6.**
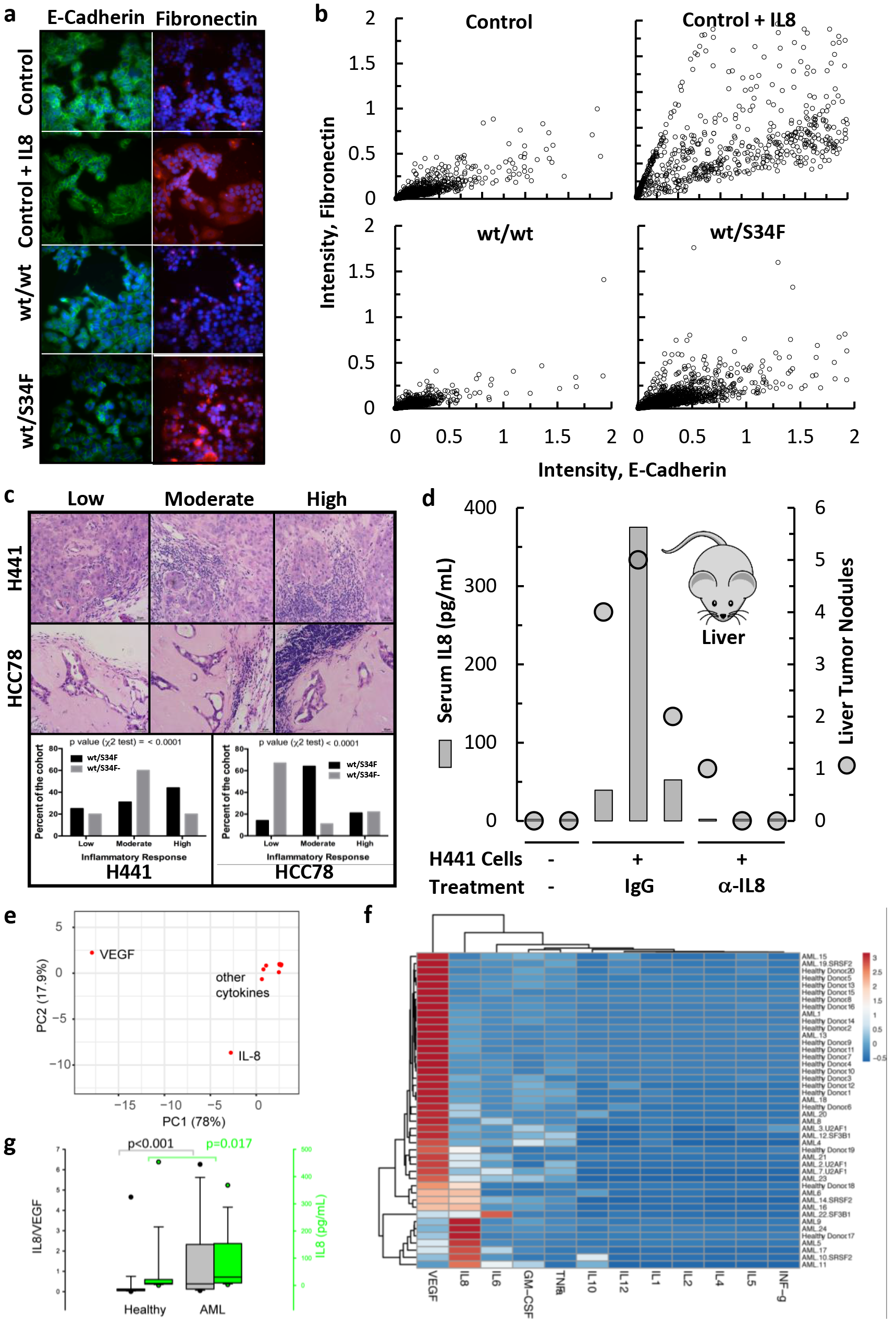
IL8 can induce EMT, inflammation and tumor progression. a. MCF-7 cells were cultured in either control, control + IL8 (1nM) or conditioned media (EMT assay) from WT or S34F mutant cells. After 10 days in culture, cells were immunostained for E-cadherin (green) and Fibronectin (red). b. Scatter plot showing EMT associated changes in E-cadherin and Fibronectin expression observed by quantitative immunofluorescence detection of E-cadherin and Fibronectin in PFA fixed individual cells from panel a. c. Representative H&E stained images (top panel) showing exent of peri- or intra-tumoral iinflammation response in xenograft tumors and its quantitation (bottom panel). Scale bar, 20 μm. The number of independent tumors: H441 (wt/S34F), n = 16; H441 (wt/S34F-), n = 20; HCC78 (wt/S34F), n = 14; HCC78 (wt/S34F-), n = 9. p-values were determined by Chi-square test. d. Serum IL8 levels (gray bar) and the number of macroscopic liver nodules (black circles) from NOG mice injected (tail vein) with H441 isogenic cells, and treated with either isotype control or anti-IL8 antibodies for 10 weeks (n = 3 for each treatment group). Two naïve NOG mice (the two samples on the left) were used as negative controls. e. Principal component analysis of 12 cytokine levels in bone marrow from healthy and relapsed or refractory acute myeloid leukemia donors (RR-AML). f. Heatmap showing the clustering of cytokine measurements. Rows are centered with unit variance scaling applied to rows. Both rows and columns are clustered using Euclidean distance and average linkage. The tree ordering for rows and columns places the higher median first. Patient and spliceosome mutational status are not part of the clustering and are indicated on the right. g. IL8/VEGF ratio and IL8 levels from healthy and AML individuals. VEGF and IL8 account for 96% of the variation within the samples.

We then examined the role of the U2AF1-S34F mutation using xenografts. We used two different human lung cancer lines (HCC78 and H441) with naturally occurring S34F mutations isolated from patients and generated derived lines with frame-shifted mutant alleles as previously reported^12^. H441 cells with or without S34F mutation were able to form subcutaneous xenograft tumors in athymic nude mice with no apparent differences in tumor sizes, while HCC78-derived cells were unable to establish tumor growth over a period of 150 days^12^. Presentation of inflammatory cells at the periphery or inside the xenograft tumors (from H441) cells or residue cells (from HCC78 cells) were scored. In both cases, the occurrence of the tumor-induced inflammation was reduced by removing the S34F mutant copy from the human cell lines (Fig. 6c).

Next, we sought to assess the specific role of IL8 in mediating the ability of H441 cancer cells to form tumors in mice. One caveat in these studies is that IL8 is not present in rodents, but homologs of the IL8 receptors (CXCR1, CXCR2) are present. NOG mice were injected with H441 cells via tail vein and were treated with neutralizing IL8 antibody or with an isotype-matched IgG three times a week for ten weeks. Serum IL8 levels and tumor burden were then assessed after the mice were sacrificed three days after the last antibody injection. Half of the mice (n=3) were given intraperitoneal injections of the anti-IL8 neutralizing antibody, and the other half were injected with the control IgG. Tumors were observed in the liver, but predominantly in the lung (Fig. 6D, Extended Data Fig. 6). Although lung tumor burden did not change (Extended Data Fig. 6), the tumor burden in the liver showed a significant decrease with a corresponding decrease in serum levels of IL8 (Fig. 6D). Taken together, studies in tissue culture and xenograft models demonstrate that EMT, immune infiltration, and tumor burden are sensitive to the U2AF1 mutational status, and these effects are consistent with and/or mediated by changes in IL8.

Finally, we asked whether IL8 is elevated in relapsed/refractory acute myeloid leukemia (RR-AML), where spliceosome mutations are common^48^. We isolated serum from 24 RR-AML patients and 20 age-matched healthy donors. A third of the AML patients had known spliceosome mutations (U2AF1, n=3; SRSF2, n=3; and SF3B1, n=2). Levels of 12 cytokines/growth factors were quantified using a Luminex assay (Extended Table 5). Principal component analysis^49^ revealed that 96% of the variation in samples could be accounted for by two factors: VEGF and IL8 (Fig. 6e). Healthy individuals showed high VEGF and low IL8 levels, while the RR-AML patients showed low VEGF and high IL8 levels (Fig. 6f). IL8 alone is significantly elevated in RR-AML patients compared to healthy donors (p=0.017, Fig. 6g), and the ratio of IL8/VEGF allowes superior discrimination (p=0.0007). On this ‘axis’ of IL8/VEGF, U2AF1 mutations cluster at an intermediate point, where VEGF is decreasing and IL8 is increasing (Fig. 6f). Thus, U2AF1 S34F is not associated with the highest levels of IL8 in human AML but may function in a general pathway whereby IL8 begins to condition the niche which supports proliferation of leukemic blasts.

## Discussion

Here we show that U2AF1 can function in a non-canonical role as a translational repressor in the cytosol. This pathway was revealed by the somatic S34F mutation which occurs in cancer, resulting in translational de-repression of hundreds of mRNA, one of which (IL8) directly contributes to the oncogenic phenotype. A simple explanation for the repressive effect is that U2AF1 binds mRNA and stabilizes a translationally repressive state. In the presence of the S34F mutant, this repression is partially relieved either by direct competition, or because the mutant selectively binds mRNA to stabilize a translationally permissive state, resulting in lower overall binding of the U2AF1 protein. Importantly, all of the phenotypes we report are rescued by frameshifting the mutant allele, arguing for a gain of function dominant negative effect rather than simple loss of function.

U2AF1 selectively targets transcripts which code for the translation machinery, and the transcripts that show the greatest de-repression in the presence of the S34F mutation are involved in protein transport. Thus, there is the potential for both direct and indirect changes to the proteome. In fact, the substantial change we see in extracellular IL8 may be due both to direct changes due to decreased U2AF1 binding and also changes in translation, processing, and trafficking of mature IL8 which are indirect effects. Translation initiation is often mis-regulated in cancer^50^ and the translation machinery is tightly controlled at the post-transcriptional level through the mTOR pathway, which is responsive to an array of environmental conditions^51^. The oncogenic phenotypes we report here are sustained viability in the presence of extreme genotoxic stress (20 Gy X-rays) or cell non-autonomous effects such as enhanced EMT and immune infiltration mediated by elevated IL8 levels even in the absence of stress. Since IL8 contributes to recruitment of myeloid cells which suppress an adaptive immune response, one prediction from this latter effect is that cells with splicing factor mutations will be refractory to immunotherapy unless treated with neutralizing IL8 antibody. However, we speculate that the U2AF1 S34F mutation could function in a cell-autonomous, pro-growth manner in tissue through this mechanism of translational mis-regulation.

## Materials and Methods

### Plasmids, cell lines and cell cultures

The plasmid expressing the IL8-2A-GFP reporters were constructed by inserting a synthetic DNA fragment coding for wt or mutant IL8-2A-GFP between *Not*I and *Sph*I sites of a lentiviral packaging vector. Transcription from this reporter is driven off an Ubiquitin promoter. Construction of WT, S34F and frameshift HBEC lines has been described previously^12^. To generate reporter cell lines, WT or S34F mutant HBEC lines^12^ were transduced with different titers of lentivirus packaged with either WT or mt 5’-UTR IL8-2A-GFP reporter constructs. Reporter RNA expression was measured by smFISH using fluorescently labeled probes (Biosearch Technologies, San Francisco, CA) against GFP and cell populations with similar RNA levels were selected and expanded for further use. All HBEC lines were cultured in Keratinocyte Serum Free Media supplemented with bovine pituitary extract and EGF as prescribed by the manufacturer (Invitrogen, Carlsbad, CA) at 37°C and 5% CO_2_.

### Irradiation and WST-1 assay

WT or S34F mutant cells (1 × 10^^4^/well) were plated in 96 well plates and grown for 24hrs before irradiation with x-rays (20Gy) (X-Rad 320, Precision X-Ray, North Branford, CT) and cultured for 6 days post irradiation. Cell viability was measured at indicated times using the WST-1 (ROCHE/SIGMA, St Louis, MO) assay measuring the conversion of tetrazolium salts to formazan by cellular tetrazolium reductase as per manufacturer’s instructions for 1hr at 37°C and 5% CO_2_.

### ELISA

HBECs were plated (1 × 10^^6^/well) in a 6 well plate and irradiated with 20Gy of X-rays. At indicated times, media was collected, filtered for cell debris using a 0.22 mm filter and snap frozen until ELISA was performed. ELISA was performed in duplicate for each sample using Quansys Human Cytokine-Inflammation 9-plex kit (Quansys Biosciences, Logan, UT) containing IL1α, IL1 β, IL2, IL4, IL6, IL8, IL10, IFNγ & TNfα. The cytokine and chemokine levels were determined from the standard curve developed with each experiment using standards provided by the manufacturer. The anylate levels (pg/mL) are the mean of three independent experiments with standard deviation.

### Immunofluorescence assay, image acquisition and analysis

Cells were seeded at ^~^30% confluency on 1.5mm thick round cover glass in 12 well plates and cultured under normal growth conditions (5% CO_2_, 37°C) up to ^~^80% confluency. Cells were washed once with PBS and fixed in 4% PFA (made in PBS) for 15 min at 22 °C. Cells were then washed thrice with PBS for 10 min each at 22 °C and permeabilized in 0.1% Triton X-100 (in PBS) for 15 min at 22 °C. Cells were washed thrice in PBS, blocked with 5% normal calf serum (in PBS) for 2h at 22 °C and incubated in primary antibody diluted in 5% normal calf serum at 4 °C overnight. Cells were washed thrice in PBS and incubated with fluorescently labeled secondary antibodies diluted in 5% normal calf serum for 1h at 22°C, washed thrice in PBS, the cover glass mounted on glass slide using ProLong Gold with DAPI (Invitrogen, Carlsbad, CA) and cured overnight at 22 °C. The slides were imaged on a custom-built epifluorescence Rapid Automated Modular Mounting (RAMM Scope; ASI, Eugene, OR) base mounted with a Plan-Apochromatic 40X (NA 1.4) objective (Zeiss, USA), illuminated with a LED light source (Model Spectra-6LCR-SA; Lumencor, Beaverton, OR) and the emitted fluorescence collected with a CMOS camera (Hamamatsu ORCA-Flash 4.0; Hamamatsu, Japan). Z-stacks (0.5 μM) were acquired to cover the entire thickness of the cells and the stacks were sum projected. Cells or nuclei were segmented and the mean fluorescence intensity calculated using the open source software, CellProfiler 2.1.1 (http://cellprofiler.org/). Source and dilutions of primary and secondary antibodies used for immunofluorescence: IL8 (eBioscience-BMS136, 1:1000); mH2A1.1 (Cell Signaling Technologies-D5F6N, 1:1500); mH2A1.2 (EMD Millipore-MABE61, 1:1000); NPM1 (AbCam-AB10530, 1:1000); IPO7 (Invitrogen-PAS21764, 1:1000); U2AF1 (AbCam-AB172614, 1:1500); U2AF2 (Santa Cruz Technologies-SC53942, 1:1000); TSC2 ThermoFisher-37-0500, 1:1000); RUNX1 (ThermoFisher-MA5-15814, 1:1000); TRIM28 (ThermoFisher-MA1-2023, 1:1000); Fibronectin (AbCam-AB23750, 1:1500); E-cadherin (Life Technologies-131700, 1:1000); Anti-Mouse IgG-Alexa 570 (Life Technologies-A11004, 1:2500); Anti-Rabbit IgG-Alexa 670 (Life Technologies-A21244, 1:2500).

### Single Molecule RNA Fluorescence In situ Hybridization (smRNA FISH)

Cells were plated at ^~^30% confluency on 1.5mm thick round cover glass in 12 well plates and cultured under normal growth conditions (5% CO_2_, 37°C) up to ^~^80% confluency. Cells were washed once with PBS, fixed in 4% PFA (made in PBS) for 15 min at 22°C and smRNA FISH performed using fluorescent probes directed against IL8 mRNA or GFP mRNA as described previously^52^. Cells and nuclei were segmented using CellProfiler 2.1.1, single molecule RNA localization in cells and transcripts/cell calculated using software custom written in IDL (http://www.harrisgeospatial.com/SoftwareTechnology/IDL) and available at www.larsonlab.net.

### RNA Immunoprecipitation

Cells were grown to ^~^90% confluency and 5 x 10^7^ cells were used for isolating cytoplasmic fraction as described previously^53^. Briefly, the cells were washed in ice cold PBS, cells suspended (1 ml buffer/50mg cells) in hypotonic lysis buffer (10 mM Tris-HCl pH 7.5, 10 mM NaCl, 3 mM MgCl_2_, 0.1% NP-40, 10% Glycerol) supplemented with protease inhibitors (ROCHE-SIGMA, St. Louis, MO) and RNASin (Promega, Madison, WI) as suggested by the manufacturer and lysed on ice for 10 min. The cell suspension was centrifuged at 800g for 8 min at 4 °C and the supernatant collected as cytoplasmic fraction. The cytoplasmic fraction was adjusted to 150mM NaCl by addition of 5M NaCl and centrifuged at 18000g for 15 min at 4°C. The supernatant was first pre-cleared with Anti-rabbit IgG conjugated Dynabeads (Invitrogen, Carlsbad, CA) for 2h at 4 °C and U2AF1 associated RNA was then immunoprecipitated from the cytoplasm (overnight at 4 °C) with anti U2AF1 antibodies (AbCam-AB172614). The immunoprecipitate was then adsorbed on Anti-rabbit IgG conjugated Dynabeads at 2 fold excess of secondary antibody bound beads relative to primary antibodies for 2h at 4 °C. The beads were washed four times with high salt buffer (50 mM Tris-HCl pH 7.5, 500 mM NaCl, 4 mM MgCl_2_ and 0.05% NP-40) followed by four washes with low salt buffer (50 mM Tris-HCl pH 7.5, 150 mM NaCl, 4 mM MgCl_2_ and 0.05% NP-40). The samples were suspended in low salt buffer and used for RNA preparation or fractionation on polyacrylamide gels for western blot analysis and probed with indicated antibodies.

### Isolation of polysomes

Polysomes were isolated from cells cultured in one 10 cm dish at ^~^80% confluency as described in^42^. The cells were incubated in media containing 10 μg/ml cycloheximide for 10 min under normal culture conditions, washed with PBS, trypsinized, washed with ice-cold PBS containing 10 μg/ml cycloheximide and lysed in lysis buffer (20 mM Tris pH 7.2, 130 mM KCl, 15 mM MgCl2, 0.5% (v/v) NP-40, 0.2 mg/mL Heparin, protease inhibitors (Sigma, St Louis, MO), RNASin (Promega, Madison, WI), 2.5 mM DTT, 0.5% deoxycholic acid and 10 μg/ml cycloheximide. The extract was clarified by centrifugation (8000g for 10 min at 4 °C). The clarified lysate was loaded onto two 10- 50% linear sucrose gradient (10 mM Tris pH 7.2, 60 mM KCl, 15 mM MgCl2, 1 mM DTT, 0.5% (v/v) NP-40, mg/mL Heparin) and centrifuged (40000 rpm for 2h at 4 °C) in an SW41 Ti rotor (Beckman, Palo Alto, CA). UV profiling (254 nm) of ribosomes and fractionation of polysomes was done using a fractionator (Biocomp Instruments, Fredericton, Canada).

### RNA Isolation and RT-PCR

RNA was isolated from immune complexes using Qiagen RNA purification kit (Qiagen, Germantown, MD) as per manufacturer’s instructions, and from polysome fractions using phenol:chloroform extraction followed by ethanol precipitation using standard protocols. 100 ng of RNA was reverse transcribed using oligo-dT primer_15_ and Protoscript II (NEB, Ipswich, MA) as per manufacturer’s instructions. qPCR of RIP samples was performed using iQ SYBR Green Supermix (BIO-RAD, Hercules, CA) using 1 μL of RT reaction product. Semi-quantitative PCR was performed using 1 μL of RT reaction product from polysome fractions for 18 cycles. The reaction products were separated on a 1.5% agarose gel and quantified using open source software ImageJ (https://imagej.nih.gov/ij/).

### EMT assay

MCF-7 cells were cultured in MEM supplemented with 5% FBS. For the EMT assay, MEM media was replaced with filtered conditioned KSFM media collected from 3-day cultures of either WT or S34F mutant HBECs and cells cultured for another 6-8 days in conditioned media, fixed and stained with antibodies against Fibronectin or E-Cadherin as described above.

### FACS

Flow cytometric analysis of GFP expression in HBECs was done using BD FACSCalibur (BDBiosciences, San Jose, CA). GFP expression from wt/wt or wt/S34F cells expressing wt- or mt-5’-UTR IL8-2A-GFP reporter were analysed using FACS. Cells were trypsinized, neutralized with Trypsin Neutralizing Reagent and washed with PBS before FACS analysis. wt/wt cells served as a control for autofluorescence and the reporter cell lines were gated above the autofluorescence of wt/wt cells (Extended Data Fig.5). Each experiment counted ^~^50K events for each cell line and the average of three independent experiments with SD presented.

### RIP- and Polysome-Seq and analysis

For each of the three cell lines used for RIP (wt/wt, wt/S34F, wt/S34F-), three samples were collected and sequenced: input, U2AF1 RIP, and IgG non-specific IP. RNA was extracted using TRIzol according to manufacturer’s instructions. rRNA was removed using the NEBNext rRNA Depletion Kit rRNA Removal Kit (NEB) followed by cDNA library preparation (NEBNext^®^ Ultra RNA Library Prep kit for Illumina). The samples were sequenced on the illumina HiSeq 3000 platform. Sequences were aligned to the human genome version hg19 using TopHat^54^. Abundances of transcript-mapped RNA sequenses were estimated using rnacounter python program. RNA isolated from individual sucrose gradient fractions were used for polysome sequencing as described above.

The following steps are used to generate a list of transcripts for analysis:

1. Only transcripts which show abundance of RPKM > 1 in all samples (except the IgG non-specific binding control, where the RPKM cutoff is 0.05) and both biological replicates are used for analysis.
2. RPKM values from the two biological replicates are averaged to generate the data in Extended Data Table 3.
3. For each cell line (wt/wt, wt/s34f, wt/-, wt/s34f-) the log_2_ ratio of the IP/input is tabulated, and for the wt/wt cell line the log_2_ ratio of IP/IgG is also included.
4. For decile analysis, transcripts are ranked either by the difference in log_2_ ratios between wt/wt and wt/s34f cells or the log_2_ ratio in wt/wt cells.

The following steps are used to compute individual and composite polysome profiles from the sequencing read count data:

1. For each sample (wt/wt, wt/s34f, wt/s34f-), 12 fractions were collected from the sucrose gradient, and the bottom/heaviest 10 fractions (Fig. 3a, fractions 3 – 12) were sequenced.
2. Fractions 10-12 were pooled to increase coverage. Fraction 5 and 6 correspond to the monosome fractions.
3. Reads mapping to particular message are normalized by the total reads in that sample and multiplied by 1×10^6^, generating counts per million (CPM) for each transcript in each fraction.
4. The CPM in each fraction constitutes the polysome profile (poly(x), where ‘x’ designates the sucrose fraction) which is now normalized for read depth across fractions/samples. This profile can be further normalized by:
  a. total area under the profile (i.e. Fig 4a) (poly(x) /[poly(3)+poly(4)+… poly(10-12)])
  b. the amount in the monomsome fraction (i.e. Fig. 4c) (poly(x)/[poly(5)+poly(6)])
5. A single “polysome/monosome ratio” is then the ratio of the heaviest fraction to the monosome fractions (i.e. Fig. 4d, f) (poly(10-12)/[poly(5)+poly(6)]).

### Splice-site finding in 5’ UTRs

Experimental 5’ UTRs reported in ref 40 were analyzed with FIMO (http://meme-suite.org/tools/fimo). Settings: motif width = 6; input sequences = 6441 sequences between 8 and 3067 bp in length. The search motif position weight matrix was: [[0.054 0.655 0.002 0.289], [1.000 0.000 0.000 0.000 0.000 0.000 1.000 0.000], [ 0.253 0.141 0.493 0.113], [ 0.241 0.189 0.199 0.371], [0.259 0.236 0.236 0.270]]

### Mouse experiments

All procedures related to mouse experiments and husbandry were approved by the Institutional Animal Care and Use Committee at the National Human Genome Research Institute (NHGRI. Protocol: G-10-6) and Weill Cornell Medicine (WCM. Protocol: 2015-017). Xenograft tumors of H441- and HCC78-derived isogenic cells were described previously^12^. Briefly, 1-5 million cancer cells were mixed with matrigel (Corning) and implanted subcutaneously in 6-8 week old nude mice (nu/nu, JAX). Tumors were harvested once they reached 10% of the body weight or up to 150 days after tumor inoculation, and were fixed in formalin. Tissue sectioning and H&E staining were done at a paid service at Histoserv. Peri- or intra-tumoral inflammation were quantified by a pathologist.

To evaluate xenograft tumor formation in mouse lung and liver, one million isogenic H441 cells (with or without S34F mutation) were mixed at equal cell number in 100 μl PBS and were injected into NOG mice (Taconic) via tail vein. Anti-IL8 and isotype control IgG1 (R & D systems) were given to the mice via intraperitoneal injection three times a week (Monday, Wednesday, Friday) starting one day after the tail vein injection for 10 weeks. Half of the mice (n=3) were given the anti-IL8 neutralizing antibody, and the other half with the control IgG. Three days after the last antibody treatment, blood was collected from live mice by submandibular bleeding and serum levels of human IL8 were measured by ELISA. Mice were then euthanized by CO2 and the macroscopic tumor nodules on the lung and liver were counted.

### Human Peripheral Blood Collection and Serum Isolation

Blood samples from relapsed or refractory acute myeloid leukemia patients and healthy donors were collected following informed consent on one of two IRB approved protocols (04-H-0012 and 07-H-0113). RR-AML patients average age was 55 (range: 22-81) and healthy donor average age was 53 (range: 24-84). Serum was extracted from serum separator tubes (BD, Franklin Lakes, NJ) per manufactor instructions and stored in aliquots at −80°C.

### Luminex Assay

The Human XL Cytokine Discovery Premixed Luminex Performance Assay Kit (FCSTM18 from R&D Systems) was used to determine levels of 12 analytes (IL8/CXCL8, IFN-gamma, IL-10, IL-2, IL-5, TNF-alpha, VEGF, IL-6, IL-4, IL-12 p70, IL-1β, and GM-CSF) in the 44 human serum samples. All serum samples were run as one batch with technical duplicates, blanks, standards, and high and low controls as per the manufacturer’s instructions. All reported values for protein concentration in pg/mL were used in the analysis, including those extrapolated beyond the range of the standards, unless the fluorescence intensity was below that of the blanks. Averaged values were used for samples run in duplicates.

## Computer code and software availability

All open source software and custom written software used in this study has been referenced in Materials and Methods with web links or provided upon request.

## Data Availability Statement

The authors declare that (the/all other) data supporting the findings of this study are available within the paper and its supplementary information files.

## Acknowledgements

The authors like to thank Dr. Harold Varmus for support, making available cell lines, reagents, other resources and for critical reading of the manuscript; Drs. Alan Hinnebusch, Sandra Wolin, Nicholas Guydosh and Jeffrey Chao for critical reading of the manuscript and valuable suggestions. MP and DRL acknowledge Dr. Heather Kalish from the Micro Analytical Immunochemistry Unit, NBIB, NIH for performing ELISA, and Katherine McKinnon and Sophia Brown from the CCR FACS Core Facility at NIH for support with FACS. FL acknowledges the support of Dr. Benjamin Durham, M.D., from MSKCC for evaluation of tumor histology; Sukanya Goswami (WCM), Danielle Miller-O’Mard (NHGRI), and members of the transgenic mouse core at NHGRI for technical assistance with the mouse experiments. This work was supported in part by the Intramural Research Programs of the National Cancer and the National Heart, Lung, and Blood Institutes of the National Institutes of Health.

## Extended Data Figure Legends

**Extended Data Figure 1.**
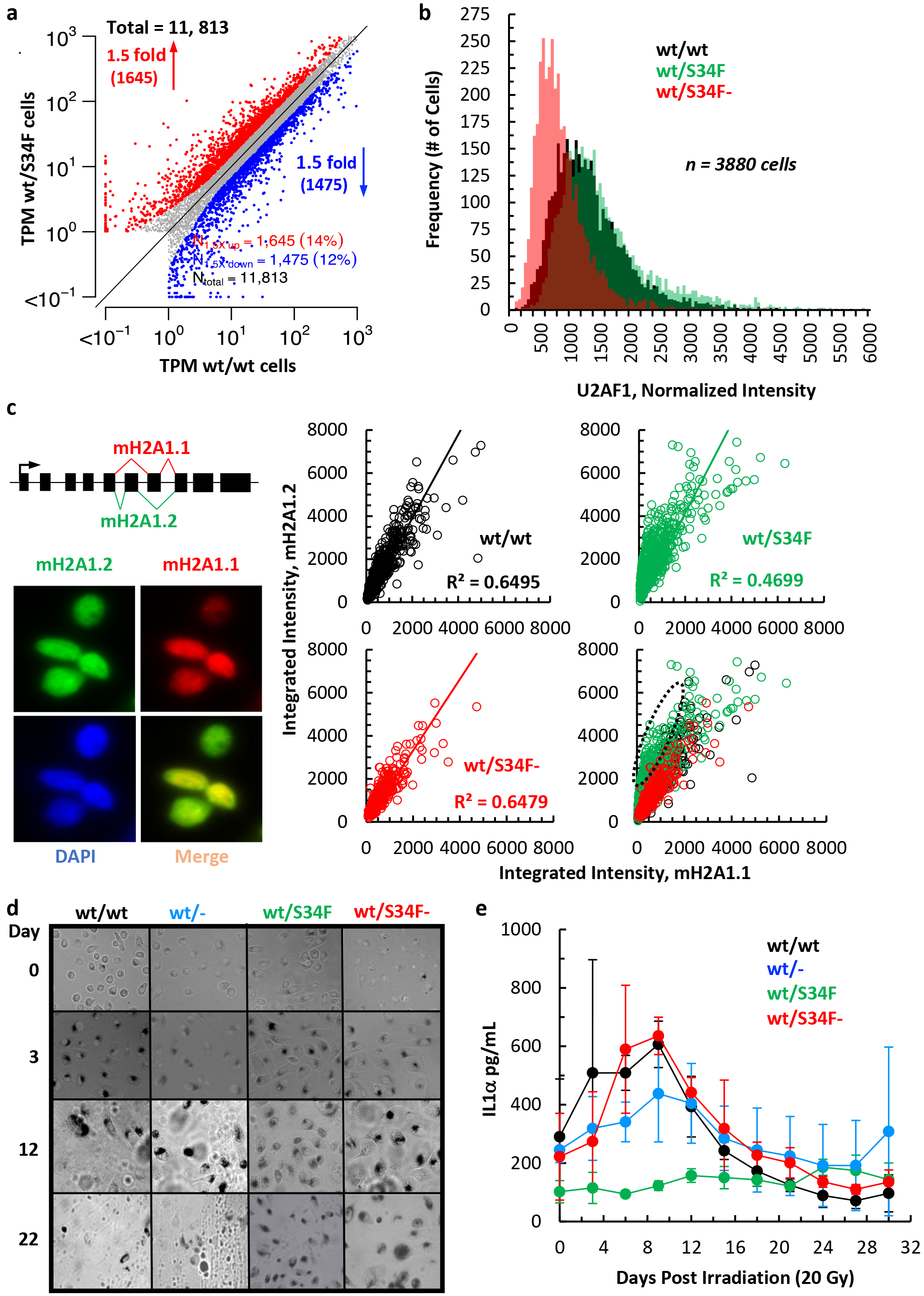
U2AF1 S34F mutant cells show altered DNA damage response and inflammatory cytokine secretion. a. Scatter plot of gene expression changes (TPM, Transcripts per million) in wt/wt and wt/S34F cells at steady state determined by RNA sequencing. The red and blue points represent genes that showed either increased (up > = 1.5 fold), or decreased (down > = 1.5 fold) expression respectively in wt/S34F cells. b. Quantitative immunostaining of U2AF1 in wt/wt, wt/S34F and wt/S34F- cells showing that U2AF1 expression is not altered in S34F mutant cells, and haploinsufficiency in the wt/S34F- cells. c. Quantitative immunostaing of macroH2A1 showing the S34F mutant induced altered splicing events results in altered protein isoform levels of macroH2A1.1 and macroH2A1.2 in the same nucleus of individual wt/wt cells. Scatter plots analysis of normalized fluorescence intensity in individual nuclei immunostained for both macroH2A1.1 and macroH2A1.2 from wt/wt, wt/S34F and wt/S34F- cells d. Representative bright field images of wt/wt, wt/-, wt/S34F and wt/S34F- cells irradiated with x-rays (20Gy), fixed and stained for β-galactosidase (stained black) after indicated times. e. Cytokine levels measured by ELISA in media collected at indicated times from wt/wt, wt/S34F, wt/- and wt/S34F- mutant cells after irradiation with x-rays (20Gy). IL1α secreted into media is presented as pg/mL (average of 3 independent experiments +/- s.d.).

**Extended Data Figure2.**
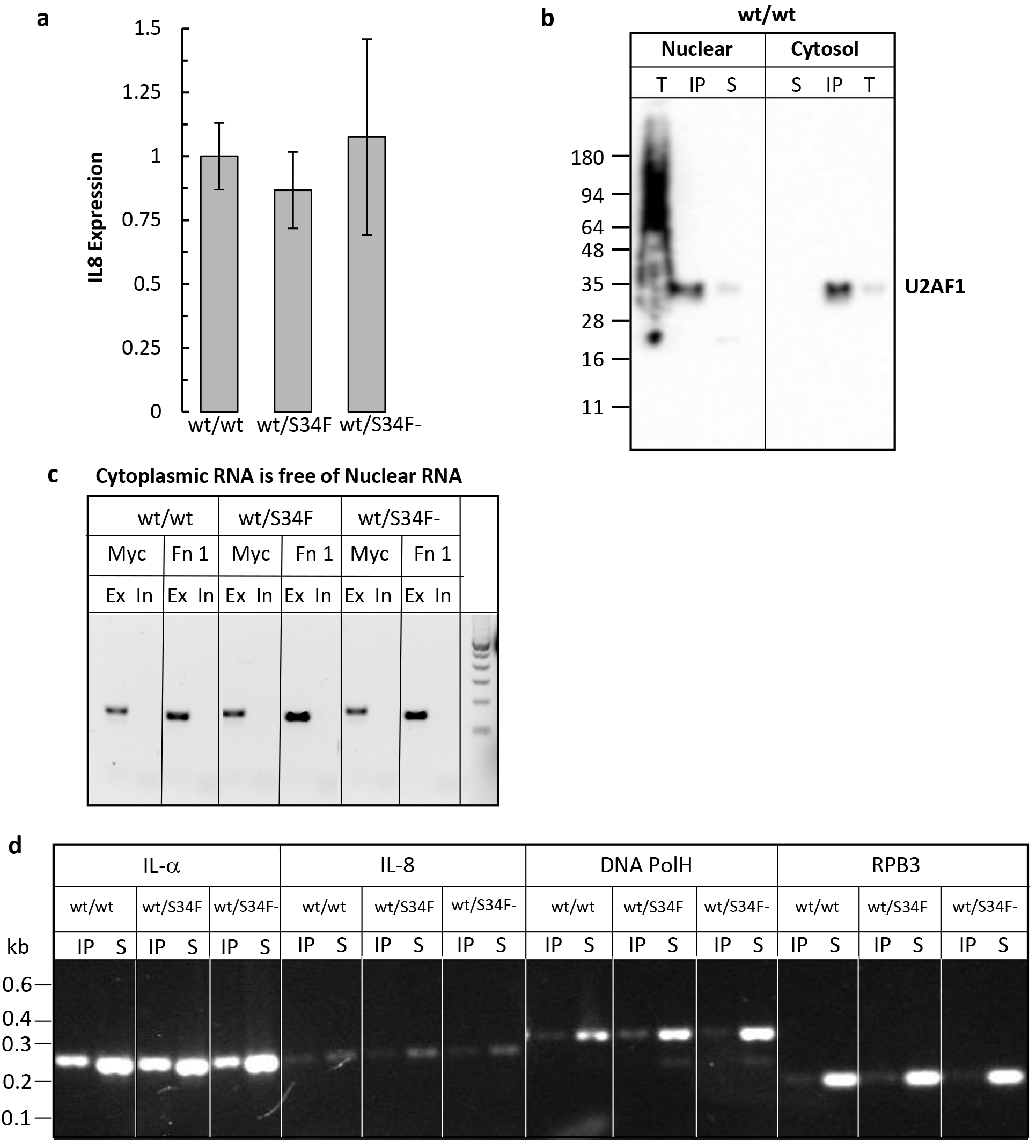
U2AF1 binds RNA in the cytoplasm. a. RT-qPCR analysis of IL8 expression in wt/wt, wt/S34F and wt/S34f- cells (3 biological replicates +/- s.d.) b. Immunoprecipitation of U2AF1 from nuclear and cytoplasmic fractions from wt/wt cells (2 fold excess of secondary antibody bound beads) followed by western blotting analysis of U2AF1 in nuclear and cytoplasmic fractions (T = total nuclear fraction, IP = immunoprecipitate and S = supernatant). c. RT-PCR analysis for detection of Myc and Fn1 pre-mRNA in cytoplasmic fraction isolated from wt/wt, wt/S34F and wt/S34F- cells (Ex = exonic primers; In = intronic primers). d. Detection of IL1α, IL8, DNA polH and RPB3 polyadenylated mRNA in U2AF1 bound cytoplasmic RIP from wt/wt, wt/S34F and wt/S34F- cells (IP = immunoprecipitate; S = IP supernatant).

**Extended Data Figure 3.**
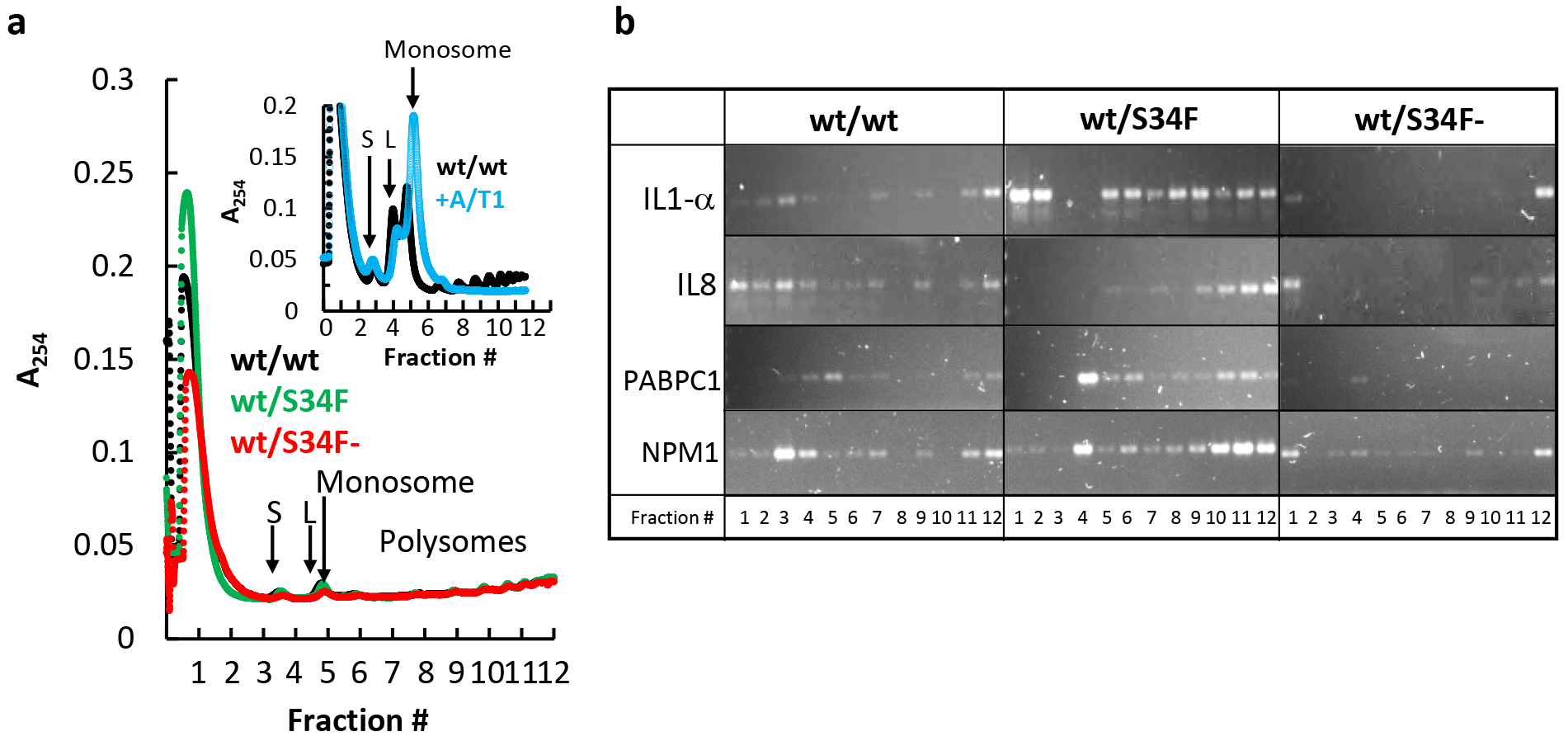
U2AF1 functions as a translational repressor in the cytoplasm. a. Polysome profiles from wt/wt (black), wt/S34F (green) and wt/S34F- (red) cells after cycloheximide treatment. Inset: profile showing collapse of polysomes into monosomes after RNAse A/T1 (blue) treatment. b. Semi-quantitative RT-PCR profiles of indicated mRNA in polysomes from wt/wt, wt/S34F and wt/S34F- cells. Limited cycle (18 cycles) PCR reaction products fractionated on 15% agarose gels.

**Extended Data Figure 4.**
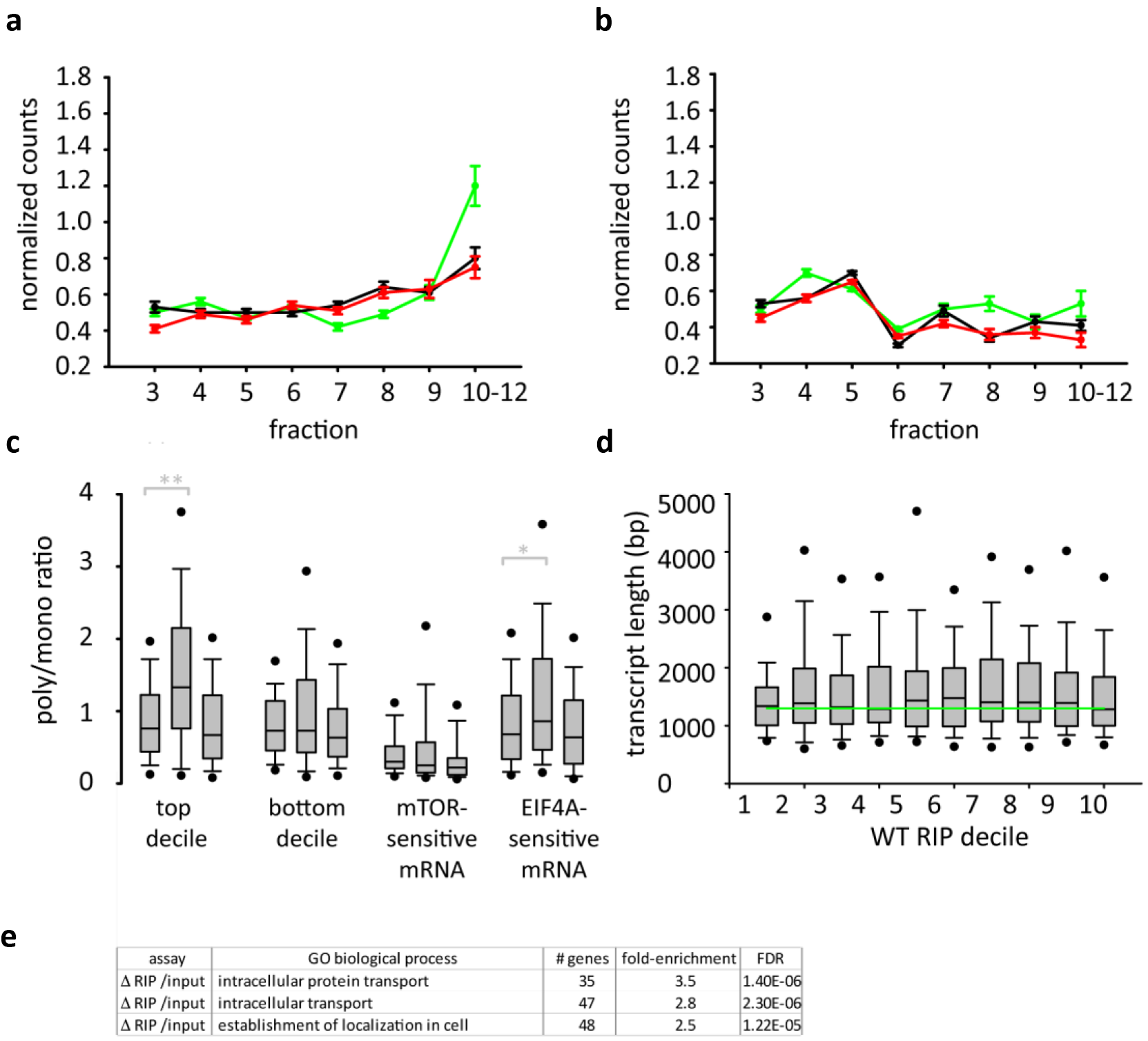
U2AF1 directly represses translation of hundreds of genes. The targets of this translational repression pathway are enriched in messages which themselves code for translation machinery, potentially resulting in direct and indirect changes in translation. EIF4A-regulated messages show increased translation in wt/S34F cells, despite not being direct binding targets. This result, however, is consistent with the change in EIF4A2 translation (Extended Data Table 3, 4). a. Average polysome profile from EIF4A-sensitive transcripts (Extended Data Table 2): polysome profiles from wt/wt (black), wt/S34F (green) and wt/S34F- (red). n=69 transcripts; bootstrap error. b. Average polysoome profile from mTOR-sensitive transcripts (Extended Data Table 2): polysome profiles from wt/wt (black), wt/S34F (green) and wt/S34F- (red). n=77 transcripts; bootstrap error. c. Polysome/monosome ratio for several different groups of transcripts. Each group contains three boxes reflecting the distribution of polysome/monosome ratios (from left to right): wt/wt, wt/S34F, wt/S34F-. The transcript groups, from left to right, are: decile 1 of changes in RIP/input from wt/wt to wt/S34F, decile 10, mTOR-sensitive transcripts, and EIF4A-sensitive transcripts. ** = p<0.001. * = p<0.05. d. mRNA length distribution within the Δ RIP deciles. There is no systematic difference in binding efficiency based on transcript length. e. GO terms for transcripts in decile 1 of changes in RIP/input from wt/wt to wt/S34F.

**Extended Data Figure 5.**
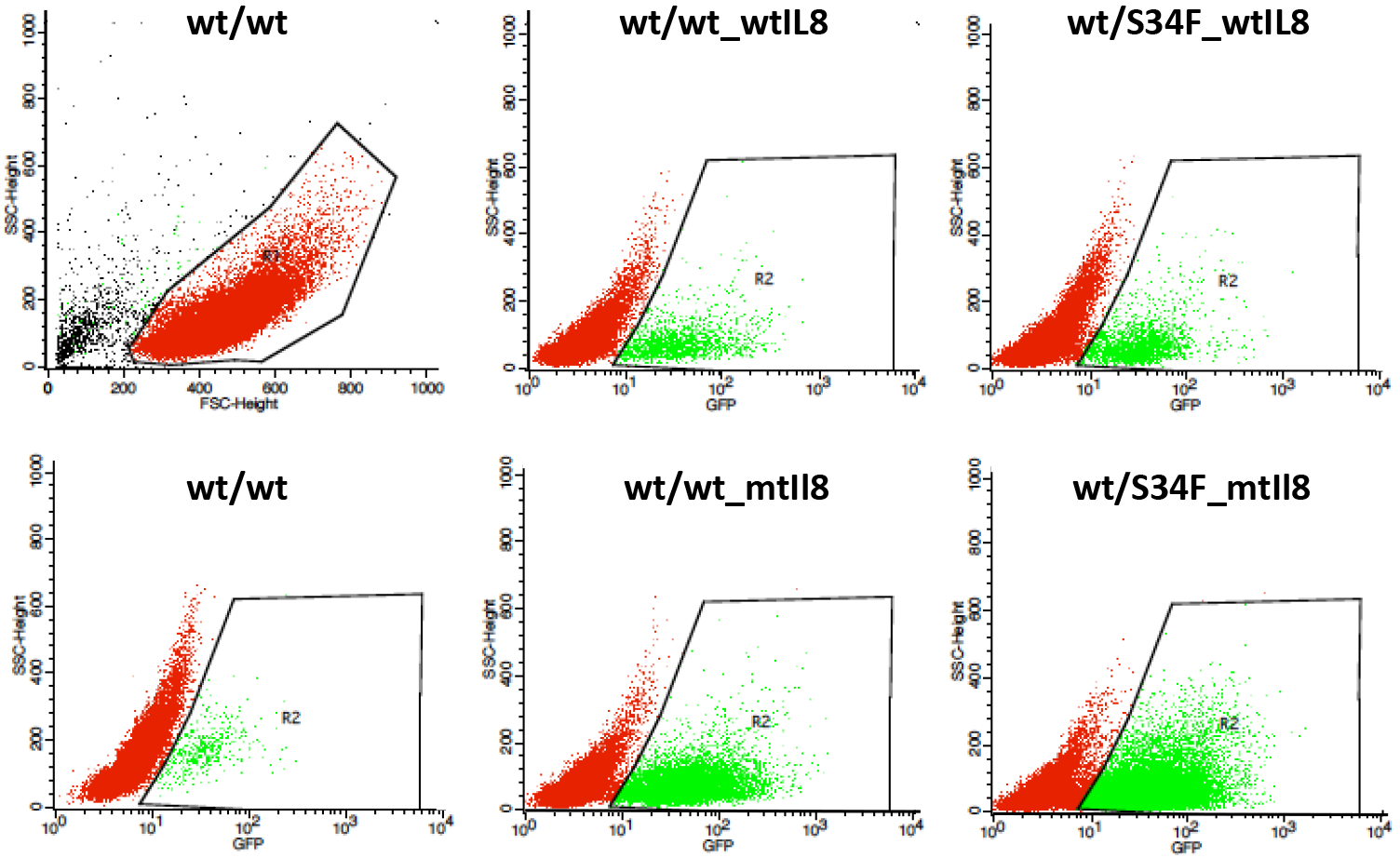
U2AF1 controls translation of IL8 via its 5’-UTR. FACS of wt/wt or wt/S34F cells expressing wt- or mt-5’-UTR IL8-2A-GFP reporter. wt/wt cells served as control for autofluorescence and the wt/wt and wt/S34F cells expressing the wt- or mt-IL8-GFP reporter were gated above the auto fluorescence from wt/wt control cells.

**Extended Data Figure 6.**
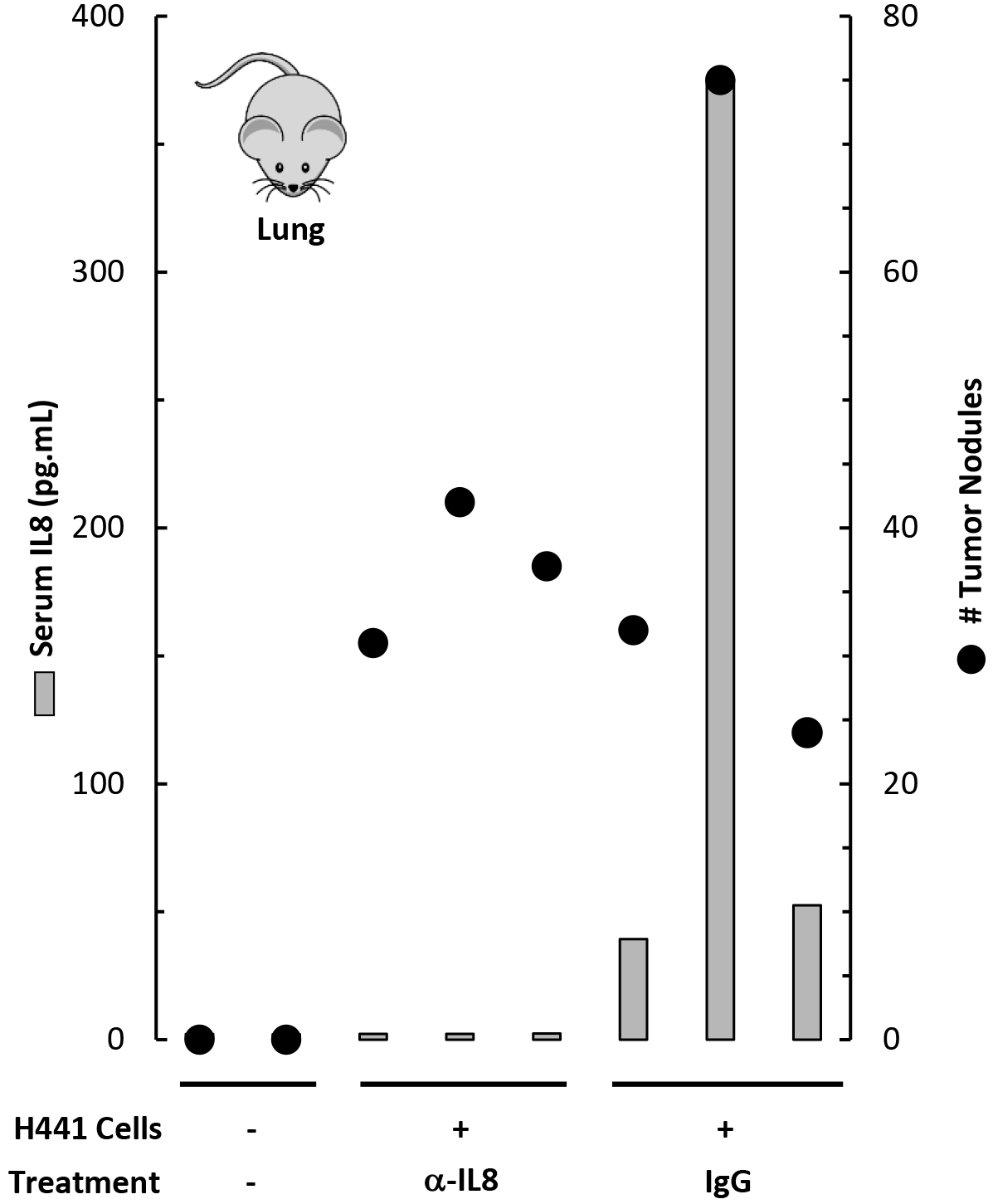
IL8 can induce EMT, inflammation and tumor progression. Serum IL8 levels (gray bar) and the number of macroscopic lung nodules (black circles) from NOG mice injected (tail vein) with H441 isogenic cells, and treated with either isotype control or anti-IL8 antibodies for 10 weeks (n = 3 for each treatment group).

**Extended Data Table 1.**
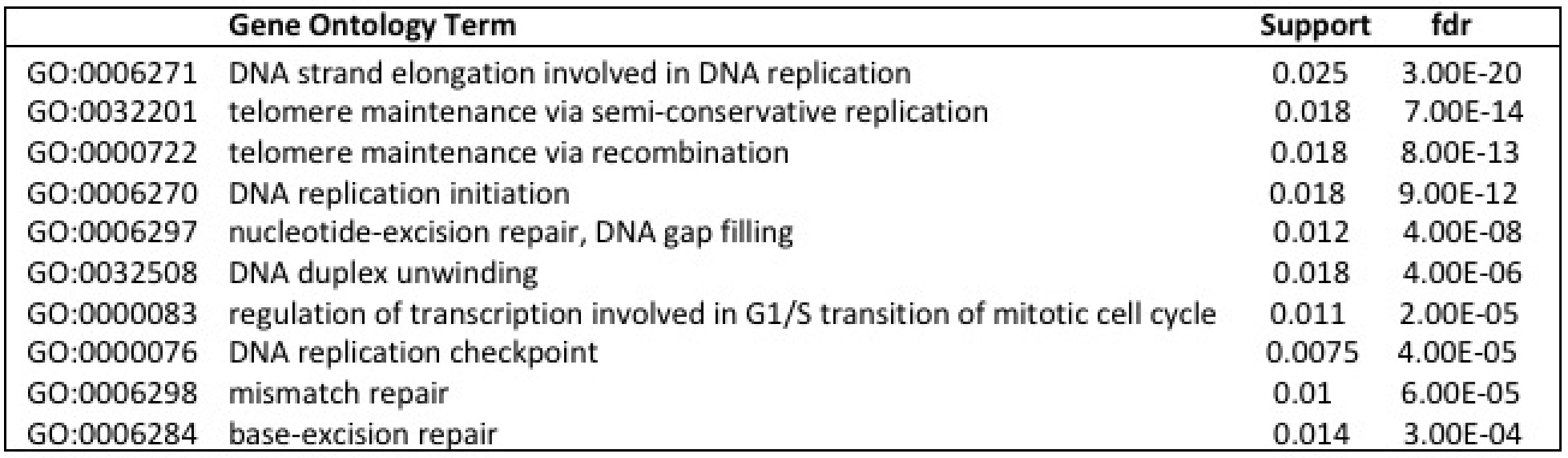

## References

1 Yoshida, K. et al. Frequent pathway mutations of splicing machinery in myelodysplasia. Nature 478, 64–69, doi:10.1038/nature10496 (2011).

2 Dvinge, H., Kim, E., Abdel-Wahab, O. & Bradley, R. K. RNA splicing factors as oncoproteins and tumour suppressors. Nat Rev Cancer 16, 413–430, doi:10.1038/nrc.2016.51 (2016).

3 Graubert, T. A. et al. Recurrent mutations in the U2AF1 splicing factor in myelodysplastic syndromes. Nat Genet 44, 53–57, doi:10.1038/ng.1031 (2011).

4 Seiler, M. et al. Somatic Mutational Landscape of Splicing Factor Genes and Their Functional Consequences across 33 Cancer Types. Cell Rep 23, 282–296 e284, doi:10.1016/j.celrep.2018.01.088 (2018).

5 Inoue, D., Bradley, R. K. & Abdel-Wahab, O. Spliceosomal gene mutations in myelodysplasia: molecular links to clonal abnormalities of hematopoiesis. Genes Dev 30, 989–1001, doi:10.1101/gad.278424.116 (2016).

6 Merendino, L., Guth, S., Bilbao, D., Martinez, C. & Valcarcel, J. Inhibition of msl-2 splicing by Sex-lethal reveals interaction between U2AF35 and the 3’ splice site AG. Nature 402, 838–841, doi:10.1038/45602 (1999).

7 Wu, S., Romfo, C. M., Nilsen, T. W. & Green, M. R. Functional recognition of the 3’ splice site AG by the splicing factor U2AF35. Nature 402, 832–835, doi:10.1038/45590 (1999).

8 Zorio, D. A. & Blumenthal, T. Both subunits of U2AF recognize the 3’ splice site in Caenorhabditis elegans. Nature 402, 835–838, doi:10.1038/45597 (1999).

9 Brooks, A. N. et al. A pan-cancer analysis of transcriptome changes associated with somatic mutations in U2AF1 reveals commonly altered splicing events. PLoS One 9, e87361, doi:10.1371/journal.pone.0087361 (2014).

10 Imielinski, M. et al. Mapping the hallmarks of lung adenocarcinoma with massively parallel sequencing. Cell 150, 1107–1120, doi:10.1016/j.cell.2012.08.029 (2012).

11 Yoshida, H. et al. A novel 3’ splice site recognition by the two zinc fingers in the U2AF small subunit. Genes Dev 29, 1649–1660, doi:10.1101/gad.267104.115 (2015).

12 Fei, D. L. et al. Wild-Type U2AF1 Antagonizes the Splicing Program Characteristic of U2AF1-Mutant Tumors and Is Required for Cell Survival. PLoS Genet 12, e1006384, doi:10.1371/journal.pgen.1006384 (2016).

13 Jenkins, J. L. & Kielkopf, C. L. Splicing Factor Mutations in Myelodysplasias: Insights from Spliceosome Structures. Trends Genet 33, 336–348, doi:10.1016/j.tig.2017.03.001 (2017).

14 Kielkopf, C. L. Insights from structures of cancer-relevant pre-mRNA splicing factors. Curr Opin Genet Dev 48, 57–66, doi:10.1016/j.gde.2017.10.008 (2017).

15 Okeyo-Owuor, T. et al. U2AF1 mutations alter sequence specificity of pre-mRNA binding and splicing. Leukemia 29, 909–917, doi:10.1038/leu.2014.303 (2015).

16 Coulon, A. et al. Kinetic competition during the transcription cycle results in stochastic RNA processing. Elife 3, doi:10.7554/eLife.03939 (2014).

17 Furney, S. J. et al. SF3B1 mutations are associated with alternative splicing in uveal melanoma. Cancer Discov 3, 1122–1129, doi:10.1158/2159-8290.CD-13-0330 (2013).

18 Ilagan, J. O. et al. U2AF1 mutations alter splice site recognition in hematological malignancies. Genome Res 25, 14–26, doi:10.1101/gr.181016.114 (2015).

19 Przychodzen, B. et al. Patterns of missplicing due to somatic U2AF1 mutations in myeloid neoplasms. Blood 122, 999–1006, doi:10.1182/blood-2013-01-480970 (2013).

20 Park, S. M. et al. U2AF35(S34F) Promotes Transformation by Directing Aberrant ATG7 Pre-mRNA 3’ End Formation. Mol Cell 62, 479–490, doi:10.1016/j.molcel.2016.04.011 (2016).

21 Chen, L. et al. The Augmented R-Loop Is a Unifying Mechanism for Myelodysplastic Syndromes Induced by High-Risk Splicing Factor Mutations. Mol Cell 69, 412–425 e416, doi:10.1016/j.molcel.2017.12.029 (2018).

22 Bavik, C. et al. The gene expression program of prostate fibroblast senescence modulates neoplastic epithelial cell proliferation through paracrine mechanisms. Cancer Res 66, 794–802, doi:10.1158/0008-5472.CAN-05-1716 (2006).

23 Coppe, J. P. et al. Senescence-associated secretory phenotypes reveal cell-nonautonomous functions of oncogenic RAS and the p53 tumor suppressor. PLoS Biol 6, 2853–2868, doi:10.1371/journal.pbio.0060301 (2008).

24 Krtolica, A., Parrinello, S., Lockett, S., Desprez, P. Y. & Campisi, J. Senescent fibroblasts promote epithelial cell growth and tumorigenesis: a link between cancer and aging. Proc Natl Acad Sci U S A 98, 12072–12077, doi:10.1073/pnas.211053698 (2001).

25 Liu, D. & Hornsby, P. J. Senescent human fibroblasts increase the early growth of xenograft tumors via matrix metalloproteinase secretion. Cancer Res 67, 3117–3126, doi:10.1158/0008-5472.CAN-06-3452 (2007).

26 Grivennikov, S. I., Greten, F. R. & Karin, M. Immunity, inflammation, and cancer. Cell 140, 883–899, doi:10.1016/j.cell.2010.01.025 (2010).

27 Taniguchi, K. & Karin, M. NF-kappaB, inflammation, immunity and cancer: coming of age. Nat Rev Immunol 18, 309–324, doi:10.1038/nri.2017.142 (2018).

28 Herranz, N. et al. mTOR regulates MAPKAPK2 translation to control the senescence-associated secretory phenotype. Nat Cell Biol 17, 1205–1217, doi:10.1038/ncb3225 (2015).

29 Laberge, R. M. et al. MTOR regulates the pro-tumorigenic senescence-associated secretory phenotype by promoting IL1A translation. Nat Cell Biol 17, 1049–1061, doi:10.1038/ncb3195 (2015).

30 Fan, J. et al. Chemokine transcripts as targets of the RNA-binding protein HuR in human airway epithelium. J Immunol 186, 2482–2494, doi:10.4049/jimmunol.0903634 (2011).

31 Park, H. Y. et al. The Arabidopsis splicing factors, AtU2AF65, AtU2AF35, and AtSF1 shuttle between nuclei and cytoplasms. Plant Cell Rep 36, 1113–1123, doi:10.1007/s00299-017-2142-z (2017).

32 Gama-Carvalho, M., Carvalho, M. P., Kehlenbach, A., Valcarcel, J. & Carmo-Fonseca, M. Nucleocytoplasmic shuttling of heterodimeric splicing factor U2AF. J Biol Chem 276, 13104–13112, doi:10.1074/jbc.M008759200 (2001).

33 Aviner, R. et al. Proteomic analysis of polyribosomes identifies splicing factors as potential regulators of translation during mitosis. Nucleic Acids Res 45, 5945–5957, doi:10.1093/nar/gkx326 (2017).

34 Maslon, M. M., Heras, S. R., Bellora, N., Eyras, E. & Caceres, J. F. The translational landscape of the splicing factor SRSF1 and its role in mitosis. Elife, e02028, doi:10.7554/eLife.02028 (2014).

35 Sanford, J. R., Gray, N. K., Beckmann, K. & Caceres, J. F. A novel role for shuttling SR proteins in mRNA translation. Genes Dev 18, 755–768, doi:10.1101/gad.286404 (2004).

36 Thoreen, C. C. et al. A unifying model for mTORC1-mediated regulation of mRNA translation. Nature 485, 109–113, doi:10.1038/nature11083 (2012).

37 Rubio, C. A. et al. Transcriptome-wide characterization of the eIF4A signature highlights plasticity in translation regulation. Genome Biol 15, 476, doi:10.1186/s13059-014-0476-1 (2014).

38 Wolfe, A. L. et al. RNA G-quadruplexes cause eIF4A-dependent oncogene translation in cancer. Nature 513, 65–70, doi:10.1038/nature13485 (2014).

39 Hsieh, A. C. et al. The translational landscape of mTOR signalling steers cancer initiation and metastasis. Nature 485, 55–61, doi:10.1038/nature10912 (2012).

40 Gandin, V. et al. nanoCAGE reveals 5’ UTR features that define specific modes of translation of functionally related MTOR-sensitive mRNAs. Genome Res 26, 636–648, doi:10.1101/gr.197566.115 (2016).

41 Abril, J. F., Castelo, R. & Guigo, R. Comparison of splice sites in mammals and chicken. Genome Res 15, 111–119, doi:10.1101/gr.3108805 (2005).

42 Panda, A. C., Martindale, J. L. & Gorospe, M. Polysome Fractionation to Analyze mRNA Distribution Profiles. Bio Protoc 7, doi:10.21769/BioProtoc.2126 (2017).

43 Fernando, R. I., Castillo, M. D., Litzinger, M., Hamilton, D. H. & Palena, C. IL-8 signaling plays a critical role in the epithelial-mesenchymal transition of human carcinoma cells. Cancer Res 71, 5296–5306, doi:10.1158/0008-5472.CAN-11-0156 (2011).

44 Ortiz-Montero, P., Londono-Vallejo, A. & Vernot, J. P. Senescence-associated IL-6 and IL-8 cytokines induce a self- and cross-reinforced senescence/inflammatory milieu strengthening tumorigenic capabilities in the MCF-7 breast cancer cell line. Cell Commun Signal 15, 17, doi:10.1186/s12964-017-0172-3 (2017).

45 Shen, T. et al. CXCL8 induces epithelial-mesenchymal transition in colon cancer cells via the PI3K/Akt/NF-kappaB signaling pathway. Oncol Rep 37, 2095–2100, doi:10.3892/or.2017.5453 (2017).

46 Thiery, J. P., Acloque, H., Huang, R. Y. & Nieto, M. A. Epithelial-mesenchymal transitions in development and disease. Cell 139, 871–890, doi:10.1016/j.cell.2009.11.007 (2009).

47 Kalluri, R. EMT: when epithelial cells decide to become mesenchymal-like cells. J Clin Invest 119, 1417–1419, doi:10.1172/JCI39675 (2009).

48 Papaemmanuil, E. et al. Genomic Classification and Prognosis in Acute Myeloid Leukemia. N Engl J Med 374, 2209–2221, doi:10.1056/NEJMoa1516192 (2016).

49 Metsalu, T. & Vilo, J. ClustVis: a web tool for visualizing clustering of multivariate data using Principal Component Analysis and heatmap. Nucleic Acids Res 43, W566–570, doi:10.1093/nar/gkv468 (2015).

50 Mamane, Y., Petroulakis, E., LeBacquer, O. & Sonenberg, N. mTOR, translation initiation and cancer. Oncogene 25, 6416–6422, doi:10.1038/sj.onc.1209888 (2006).

51 Laplante, M. & Sabatini, D. M. mTOR signaling in growth control and disease. Cell 149, 274–293, doi:10.1016/j.cell.2012.03.017 (2012).

52 Palangat, M. & Larson, D. R. Single-gene dual-color reporter cell line to analyze RNA synthesis in vivo. Methods 103, 77–85, doi:10.1016/j.ymeth.2016.04.009 (2016).

53 Gagnon, K. T., Li, L., Janowski, B. A. & Corey, D. R. Analysis of nuclear RNA interference in human cells by subcellular fractionation and Argonaute loading. Nat Protoc 9, 2045–2060, doi:10.1038/nprot.2014.135 (2014).

54 Trapnell, C. et al. Differential gene and transcript expression analysis of RNA-seq experiments with TopHat and Cufflinks. Nat Protoc 7, 562–578, doi:10.1038/nprot.2012.016 (2012).

